# Integrating single-cell multimodal epigenomic data using 1D-convolutional neural networks

**DOI:** 10.1101/2024.02.16.580655

**Authors:** Chao Gao, Joshua D. Welch

## Abstract

Recent experimental developments enable single-cell multimodal epigenomic profiling, which measures multiple histone modifications and chromatin accessibility within the same cell. Such parallel measurements provide exciting new opportunities to investigate how epigenomic modalities vary together across cell types and states. A pivotal step in using this type of data is integrating the epigenomic modalities to learn a unified representation of each cell, but existing approaches are not designed to model the unique nature of this data type. Our key insight is to model single-cell multimodal epigenome data as a multi-channel sequential signal. Based on this insight, we developed `ConvNet-VAE`s, a novel framework that uses 1D-convolutional variational autoencoders (VAEs) for single-cell multimodal epigenomic data integration. We evaluated `ConvNet-VAE`s on nano-CT and scNTT-seq data generated from juvenile mouse brain and human bone marrow. We found that `ConvNet-VAE`s can perform dimension reduction and batch correction better than previous architectures while using significantly fewer parameters. Furthermore, the performance gap between convolutional and fully-connected architectures increases with the number of modalities, and deeper convolutional architectures can increase performance while performance degrades for deeper fully-connected architectures. Our results indicate that convolutional autoencoders are a promising method for integrating current and future single-cell multimodal epigenomic datasets.

## 1 Introduction

Single-cell sequencing technologies have revolutionized our understanding of cellular heterogeneity and the complexity of biological systems. Recently, single-cell multimodal chromatin profiling has emerged as an exciting new experimental approach to investigate the cellular epigenetic landscape. Two independent studies fused nanobodies (nb) to a transposase enzyme (Tn5) and used these nb-Tn5 conjugates to detect up to three epigenome layers (histone modification or chromatin accessibility) within the same cell [1, 2]. The nano-CT and scNTT-seq technology can in principle be used to detect transcription factor binding as well, though this has not yet been demonstrated. These multimodal datasets provide simultaneous measurements of multiple epigenomic layers within individual cells, offering unprecedented opportunities to unravel how histone modifications and chromatin states drive cellular diversity. For example, one can use this type of data to investigate how different histone modifications at the same genomic locus combine to activate or repress transcription of nearby genes. However, the structure of single-cell multimodal epigenomic data is unique compared to other single-cell data types: each modality is a one-dimensional genomic track, and the total measurement for a cell consists of multiple one-dimensional tracks measured at the same genomic positions. This is quite different from any other type of single-cell data–such as scRNA-seq, snATAC-seq, CITE-seq, or 10X multiome–in which the space of features is most naturally represented in terms of genes or discrete peaks.

A number of computational approaches have been designed to perform joint dimension reduction on single-cell multimodal data types such as CITE-seq and 10X multiome. For example, the `Seurat` weighted nearest neighbor algorithm, the multi-omic factor analysis (`MOFA+`), and the `multiVI` perform linear or non-linear dimension reduction on single-cell multimodal datasets that can be represented as genes and peaks [3–5]. Approaches based on variational autoencoders (VAEs) are especially powerful for learning joint representations from single-cell multimodal data. VAEs are unsupervised probabilistic deep learning models that excel at distilling compact and meaningful representations of complex data, as evidenced by their successful applications in single-cell RNA-sequencing (scRNA-seq) data integration [6]. For multimodal problems, a VAE based on the concept of the Product of Experts (PoE) was introduced [7]. This method factorizes the joint distribution over the latent variables into a product of conditional distributions, each representing the output of a modality-specific “expert” model. Each expert is comprised of an encoder and a decoder, designed to model a specific data modality. Beyond their initial applications in image transformation and machine translation, such VAEs have been adapted for multimodal single-cell sequencing data. For example, `Cobolt` and `multiVI` use multimodal VAEs to integrate paired measurements, such as gene expression and chromatin accessibility (peaks), and learn a unified cell embedding for cell clustering and visualization [5, 8]. Although multimodal VAEs can in principle use any type of neural network layers, single-cell multimodal VAEs have only used fully-connected layers due to the unordered nature of gene features. Thus, we refer to these previous approaches as `FC-VAE`s.

However, directly applying such approaches to single-cell multimodal epigenomic data has several disadvantages. First, it requires calling peaks separately on each epigenomic layer, which results in extremely high-dimensional data because each epigenomic modality is measured across the whole genome. The number of peaks per modality usually exceeds 10^5^, and the peaks often do not overlap across modalities, further increasing the number of peaks as the number of modalities per cell increases. Second, by using a peak-centric feature representation, previous approaches neglect the ordered sequential nature of single-cell epigenomic data, in which the epigenomic state of a particular locus shares strong conditional dependence with the states of loci immediately before and after it in linear genome order. Finally, using genes and peaks neglects the multi-track nature of single-cell multimodal epigenomic data, removing the crucial information of shared genome position across modalities. This third limitation is especially problematic because it prevents integration algorithms from learning the relationship among different epigenome modalities at a given position within a single cell, which is one of the key motivations for performing single-cell multimodal epigenomic measurement in the first place.

One-dimensional convolutional neural networks (1D-CNNs) have shown success in the analysis of sequential data, especially when the spatial or temporal relationships within the data are crucial [9]. In particular, deep learning models using 1D-CNN layers have been widely used in the analysis of bulk RNA-seq and bulk epigenome data. Such networks have been trained on bulk data from cell lines and tissues to predict transcriptional and epigenetic profiles from DNA sequence [10, 11]. Recently, Yuan et al. extended this line of work to single-cell ATAC-seq data: `scBasset` [12] takes DNA sequences as input and utilizes CNNs to predict chromatin accessibility in single cells. However, to our knowledge, only `FC-VAE`s have been used to perform dimension reduction and integration of single-cell data.

Here, we present a novel 1D convolutional variational autoencoder framework (`ConvNet-VAE`s) tailored for integrating single-cell multimodal epigenomic data. We model single-cell multimodal epigenomic data as a multi-channel sequential signal. A key innovation of our method is that, by performing convolution over ordered feature space, it adopts a more appropriate inference bias than VAEs with only the fully-connected layers that are suitable for unordered features. Our approach combines two streams of work: 1D CNNs for bulk genomic data and VAEs for dimension reduction of single-cell data. Importantly, our method is fundamentally different from this previous work in several key aspects: (1) we utilize a window-based genome binning strategy on the multimodal profiles from single cells and model the fragment count in each bin; (2) we use 1D convolutional layers that operate over different epigenetic modalities instead of nucleotide bases; and (3) unlike the previous multimodal VAEs, `ConvNet-VAE`s consists of only one encoder-decoder pair. We show that `ConvNet-VAE`s can leverage the strengths of both VAEs and convolution. They effectively reduce data dimensionality and extract local genomic features with a more economical parameter usage compared to that of `FC-VAE`s.

## 2 Materials and Methods

### 2.1 Generative probabilistic model of epigenomic data

We modeled the count data of a given feature (e.g., a histone modification such as H3K27ac) by using a Poisson distribution. Consider multimodal single-cell data comprised of *M* modalities from *B* different experimental batches, with a total of *N* cells. All modalities share the same set of features *G* (e.g., binned genomic regions). We represent cell *i* with a latent factor **z**^*i*^ sampled from *𝒩* (**0, I**), characterized by batch information **b**^*i*^, and a modality-specific library size factor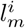. We model the generative process of the count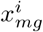 of the molecular feature *g* within modality *m* (*m ∈ {*1, 2, …, *M*}) as follows:

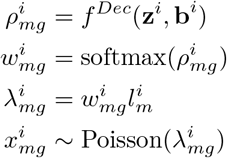

Here, 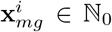 represents the count data, **z**^*i*^ *∈* ℝ ^*D*^ is the latent representation of each cell in a *D*-dimensional space, with *D* selected according to the complexity of the data. The modality-specific library size factor is denoted as 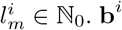is a *B*-dimensional one-hot encoded vector containing batch information.

The function *f* ^*Dec*^ denotes the decoder neural network, which consists of convolutional layers and/or fully connected layers. Through the application of a softmax activation function in the final layer, the decoder network maps the latent factors and batch label of cell *i* to the original feature space. In this study, we also implemented negative binomial (NB) distribution in the models, which is able to accommodate overdispersion in the data by including an extra parameter for dispersion. In our experiments spanning three single-cell multimodal datasets, we observed that `ConvNet-VAE`s employing negative binomial distribution exhibited negligible differences compared to those utilizing Poisson distributions (Supplementary Figure S2). Under this distributional assumption, `ConvNet-VAE`s also maintain their edge over `FC-VAE`s (Supplementary Figures S3, S4, S5). With these observations, our studies focus on Poisson-based modeling in the rest of this report.

### 2.3 Multimodal variational autoencdoers

#### Variational autoencoders (VAEs)

As previously described, we consider the observed feature vector **x**_*m*_ of a cell derived from hidden variable **z**, from batch **b**. Researchers have harnessed the VAE framework for efficient approximation of the posterior distribution for **z** [13]. VAEs, as deep generative models, exploit neural networks for variational inference, facilitating representation learning from high-dimensional data. The functionality is crucial for single-cell data integration and subsequent cell type identification [6]. Typically, VAEs are trained to optimize the evidence lower bound (ELBO) using stochastic gradient methods. In a unimodal scenario where *M* = 1, the ELBO for a feature vector **x**_1_ is defined as follows:

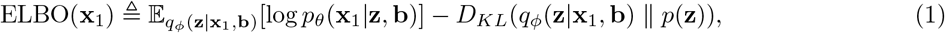

where *q*_*ϕ*_(**z**|**x**_1_, **b**) and *p*_*θ*_(**x**_*i*_|**z, b**)*p*(**z**) are the inference model (parameterized by *ϕ*) and generative model (parameterized by *θ*) respectively. To address the challenges in modeling multimodal single-cell data, we introduce multimodal VAEs in the next sections.

#### Convolutional variational autoencoders with 1D-convolutional layers (ConvNet-VAE)

Convolutional neural networks (CNNs) effectively perform tasks such as data compression and classification by learning representations of the input (for example, 1D for signals or sequences, 2D for images) [14]. In the context of 1D-CNNs, `Conv1D` filters work on the 1D input sequences and move in one direction. We introduce `ConvNet-VAE`, a variational autoencoder architecture that utilizes 1D-convolutional layers to model and integrate single-cell multimodal epigenomic data. By incorporating `Conv1D` layers, `ConvNet-VAE` efficiently embeds high-dimensional multimodal epigenomic features of the cells into a low-dimensional space suitable for clustering tasks. For compatibility with 1D-CNN, we treat the fragment count of different modalities along the binned genome as 1D sequence with multiple channels, where each channel corresponds to a different modality. Given *N* cells, then we have 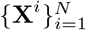, and 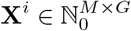 For instance, in a bi-modal setting,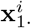 denotes the first channel, and 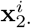 the second. The ELBO is formulated as below.

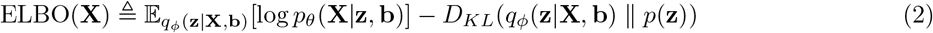

Note that we assume different modalities **x**_*m*_ (channels of **X**) are conditionally independent on **z** and **b** for tasks involving multiple modalities.

The architecture of `ConvNet-VAE` is depicted in Figure 1. This research focuses on two main configurations of `ConvNet-VAE` models. The first group of models comprises a single convolutional layer with varying sizes of kernel (K) and stride (S). The second group features multiple convolutional layers with constant kernel size and stride. By experimenting single-Conv1D-layer `ConvNet-VAE` (with a kernel size of 31 and stride of S31) on the bi-modal juvenile mouse brain dataset, we notice an increase in the marginal log-likelihood of validation data when more kernels (output channels) are applied. However, there is a disproportionately large increase in computational time compared to the gains in capturing the data distribution when the kernel count is doubled from 32 to 64 (Figure 2b). Therefore, for single-Conv1D-layer `ConvNet-VAE`s, we set the kernel count to 32. In the case of models incorporating a second or third convolutional layer, the output channels are set to 64 and 128, as is commonly done in CNN architectures. Complete specifications are provided in Table 1, including the number of feature channels produced by the convolutional layers (indicated in the parentheses). The final Conv1D layer in the decoder produces an output with a channel count that matches the number of data modalities.

**Table 1.** ConvNet-VAE Model Architecture. Two groups of `ConvNet-VAE`s are employed in this study. In Group 1, the encoders and decoders of `ConvNet-VAE`s comprise only one 1D-convolutional layer, with increasing size of the kernel (K) and stride (S) of filters (from 11 to 51). In the second group, the `ConvNet-VAE`s contain an increasing number of 1D-convolutional layers (from 1 to 3), with fixed kernel size and stride.

| ConvNet-VAE Model Architecture |  |  |  |  |  |  |  |
| --- | --- | --- | --- | --- | --- | --- | --- |
| Grp. | Kernel(K) | Stride(S) | Encoder Layers |  |  | Decoder Layers |  |
| 1 | 11 | 11 | Conv1D(32) | FC | FC($\mu, \sigma$) | FC | Conv1D |
| | 31 | 31 | Conv1D(32) | FC | FC($\mu, \sigma$) | FC | Conv1D |
| | 51 | 51 | Conv1D(32) | FC | FC($\mu, \sigma$) | FC | Conv1D |
| 2 | 11 | 3 | Conv1D(32) | FC | FC($\mu, \sigma$) | FC | Conv1D |
| | 11 | 3 | Conv1D(32) Conv1D(64) | FC | FC($\mu, \sigma$) | FC | Conv1D(32) Conv1D |
| | 11 | 3 | Conv1D(32) Conv1D(64) Conv1D(128) | FC | FC($\mu, \sigma$) | FC | Conv1D(64) Conv1D(32) Conv1D |

**Fig. 1.**
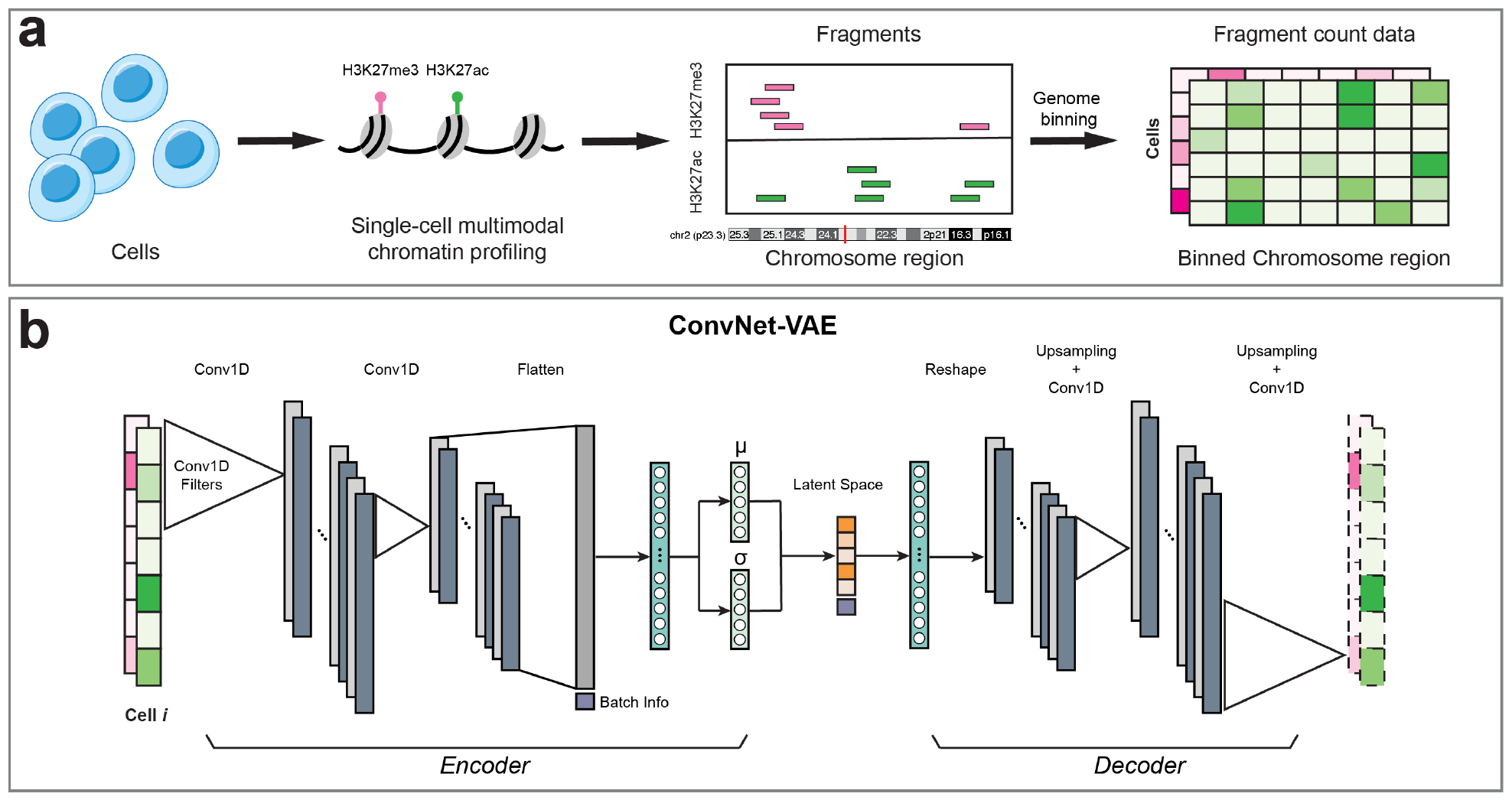
Overview of ConvNet-VAE. (**a**) For each cell, the fragments of each measured epigenomic modality are acquired by multimodal single-cell epigenome profiling (e.g., H3K27ac + H3K27me3). Followed by genome binning, we obtain the fragment count data with dimension cell *×* modality *×* Bin. (**b**) `ConvNet-VAE` applies 1D-convolution and learns low-dimensional representations of the cells from the binned multimodal fragment count of input.

**Fig. 2.**
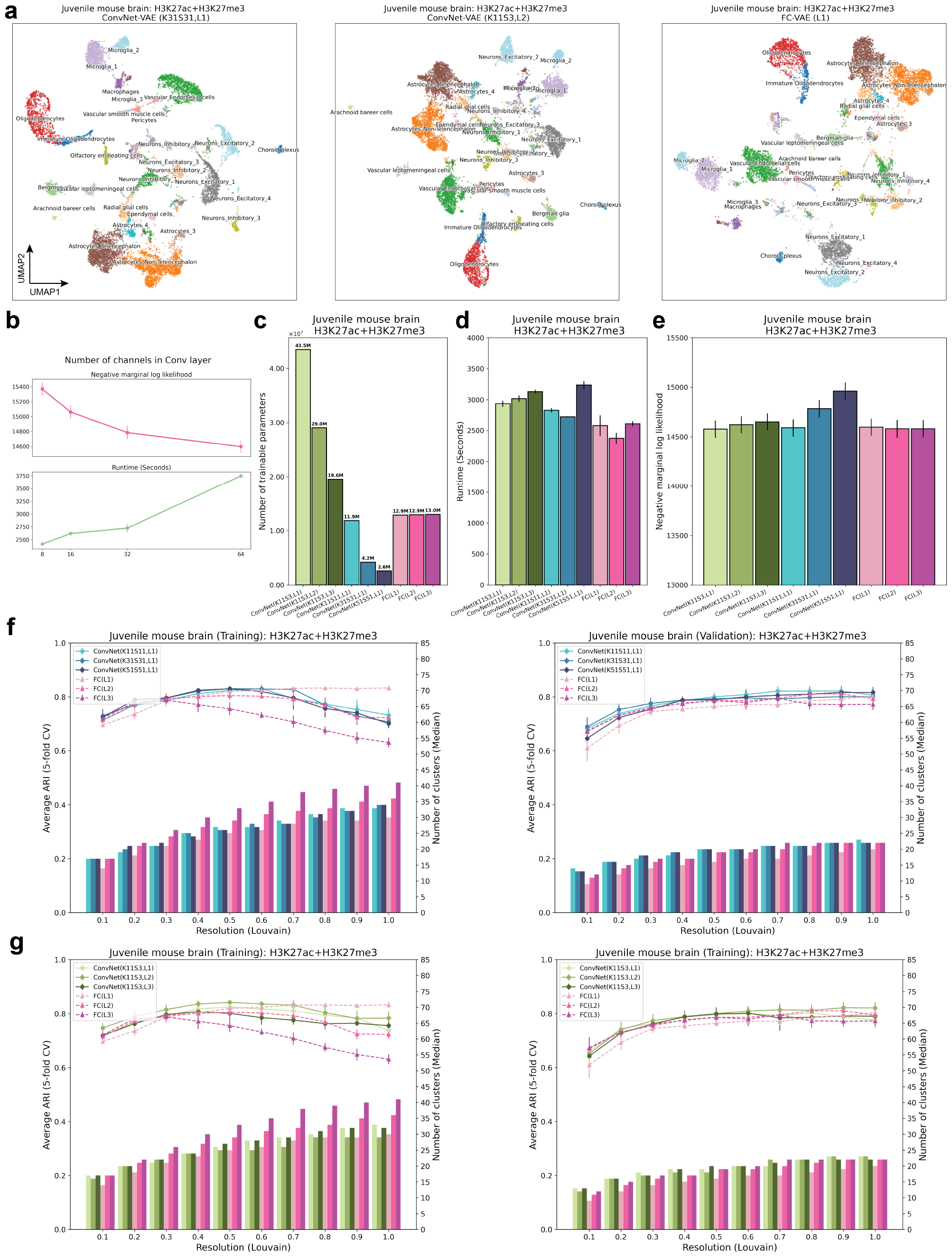
ConvNet-VAEs integrate single-cell bi-modal epigenomic profiling data from mouse brain. (**a**) UMAP visualization of cell embeddings from `ConvNet-VAE`s (Left, Middle) and `FC-VAE` (Right). (**b**) For single-Conv1D-layer `ConvNet-VAE`, more channels in the convolutional layer lead to a larger marginal log likelihood of the validation set at the cost of longer runtime in training, according to the result from 5-fold cross-validation. (**c**) The number of trainable parameters depends on the number of Conv1D layers and stride. `ConvNet-VAE`s from Group 1 (Blue) require fewer parameters than `FC-VAE`s (Pink) while those from Group 2 (Green) need more parameters. (**d**) Average training time is reported for each model. Error bars indicate standard deviation across 5-fold cross-validation. (**e**) Average negative marginal log likelihood of validation set estimated through importance sampling. A lower value implies a larger marginal log likelihood. Error bars indicate standard deviation across 5-fold cross-validation. (**f**) Comparisons between `ConvNet-VAE`s with single Conv1D layer (Group 1) and `FC-VAE`s in terms of the quality of cell embeddings (training set: left; validation set: right). The bars show the median number of clusters obtained by the Louvain algorithm from 5 splits in cross-validation over a range of resolutions. The corresponding average Adjust Rand Index (ARI) is calculated by comparing the resulting clusters to the published cell type labels, displayed as a line plot. Error bars indicate standard deviation across 5-fold cross-validation. (**g**) Comparisons between `ConvNet-VAE`s with multiple Conv1D layer (Group 2) and `FC-VAE`s in terms of cell embeddings’ quality (training set: Left; validation set: Right), exhibited in the same way as (**f**).

In general, Conv1D and FC layers are followed by Batch Normalization (1D), ReLU activation, and Dropout layers. We perform softmax activation on the output from the last decoding Conv1D layer, without any other transformation. FC(*μ,σ*) as well as the FC layer in the decoder are linear layers. The pooling layer is replaced by applying a large stride (*≥* 3). For enhanced numerical stability, each input channel—representing a different modality—undergoes log transformation (log (**x** + **1**)).

#### Variational autoencoders with fully connected layers (FC-VAE)

In order to demonstrate the advantage of `ConvNet-VAE`, we include `FC-VAE` for benchmark analyses. To address the problem of learning joint representations of multiple modalities, the idea of product-of-experts (PoE) was introduced by Wu and Goodman in 2018 [7]. We adapted the PoE approach for our specific task of multimodal inference within the context of single-cell epigenomics. Consistent with the settings described in the previous sections, we establish the following joint posterior by assuming conditional independence between *p*(**x**_*m*_|**z, b**),

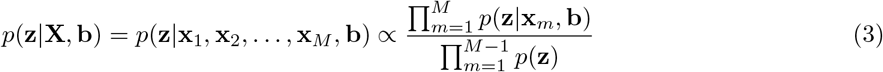

We further approximate the true single-modality posterior *p*(**z**|**x**_*m*_, **b**) using *q*(**z**|**x**_*m*_, **b**) (a Gaussian “expert”) learned from modality-specific neural networks (parameterized by *ϕ*_*m*_).

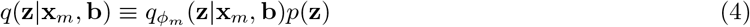

The product of Gaussian experts is still Gaussian distributed [15]. Assuming that the *m*-th expert outputs *μ*_*m*_ and *V*_*m*_ and setting 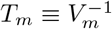 we can define the product Gaussian of **z** with the following parameters:

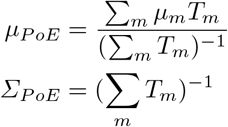

We configured `FC-VAE`s under three different settings with their architectures detailed in Table 2. Each expert model is comprised of two fully connected (FC) layers in the encoder and an additional two FC layers in the decoder, similar to that employed in `scVI` [6]. An example model architecture is shown in Figure S1. Like `ConvNet-VAE`s, `FC-VAE`s apply Batch Normalization (1D), ReLU activation, and Dropout layers in the FC layers, except for FC(*μ,σ*) and the final FC decoding layer. The models are trained to optimize the ELBO defined as 5. In addition to the comparable architectures of `FC-VAE`s and `ConvNet-VAE`s, we use exactly the same dimension (*D*) for the latent space, dropout rate, training/validation data splits, training scheme, and parameters for clustering, to ensure fair comparison (detailed in 2.7).

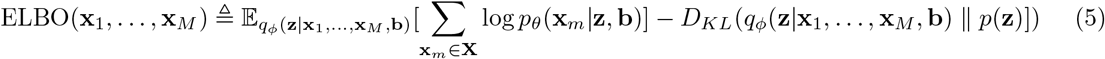

**Table 2.**
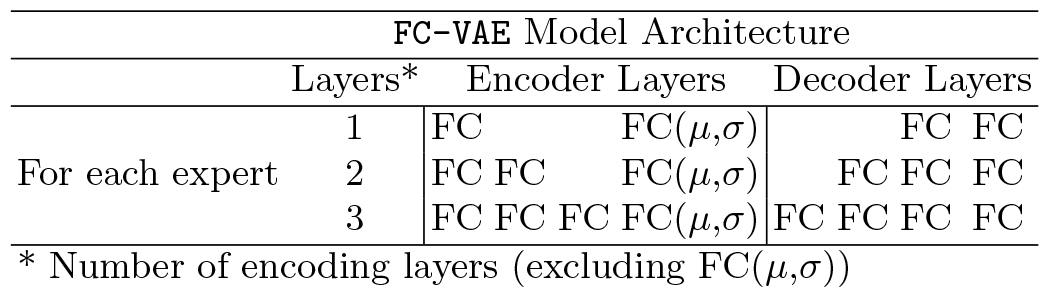
FC-VAE Model Architecture. For `FC-VAE`s based on Product of Experts, the number of layers in each expert (modality-specific encoder-decoder) differ across each `FC-VAE` variant. Taking `FC-VAE` (L1) as an example, for each expert, the encoder contains one fully connected layer in addition to one that outputs mean and log-variance parameters, and the decoder consists of two fully connected layers.

### 2.3 Evaluation on batch-effect correction

To evaluate model efficacy in data integration, we utilized four distinct metrics: ASW (Batch), graph connectivity, graph iLISI, and kBET. These metrics, introduced by Luecken et al., are tailored to assess batch-effect removal [16].

Average silhouette width (ASW) quantifies the separation of clusters. ASW (Batch) measures batch mixing, ranging from 0 to 1, where 1 indicates perfect mixing. Graph connectivity investigates how well the cells with the same identity are connected in the *k*-nearest neighbor (*k*NN) graph built from integrated data. A graph connectivity score of 1 implies good integration, where all cells with the same label are connected in the *k*NN graph. Korsunsky et al. employed integration Local Inverse Simpson’s Index (iLISI) to measure the batch distribution using local neighbors chosen on a pre-defined perplexity [17]. As an extension, Graph iLISI can take graph-based integration outputs and a higher score represents better data integration. *k*-nearest neighbor batch-effect test, known as kBET, starts by constructing a *k*NN graph, and then examines the batch label distribution in the cell’s neighborhood against the global batch label distribution through random sampling [18]. The detailed descriptions of these metrics are available in their original publications. In this benchmark analysis, we trained VAE models using the entire dataset and 5 different random initializations. We computed these metrics with default settings using the resulting cell embeddings. All metrics reported in this study are average scores across 5 runs.

### 2.4 Evaluation of VAEs’ ability to capture data distribution

To benchmark Bayesian probabilistic models in a uni-modal setting (**x**_1_), a popular strategy is to compare the marginal likelihood. A VAE model that is better at capturing the data distribution and generating samples is expected to achieve a higher marginal log-likelihood log *p*(**x**_1_) on the test set. Similarly, here we used joint conditional log-likelihood log *p*(**x**_1_, **x**_2_) and log *p*(**x**_1_, **x**_2_, **x**_3_) as the evaluation metrics in the multi-modal settings, to compare the quality of the tested deep generative models. These marginal log-likelihoods (marginal with respect to latent variable **z**) can be approximated through importance sampling [7, 19]. Assuming test data **x**_1_, **x**_2_, as well as the latent representation **z** from a given sample *i* in the bi-modal setting, hence we have

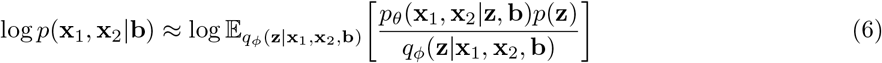

The RHS of 6 can be estimated by Eq. (7):

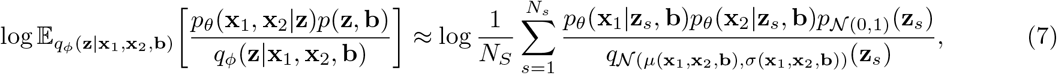

where the samples *z*_*s*_ are randomly drawn from the importance distribution *𝒩* (*μ*(**x**_1_, **x**_2_, **b**), *σ*(**x**_1_, **x**_2_, **b**)) defined by the output from the inference networks. *N*_*s*_ is the number of importance samples. *p*_*θ*_(*x*_1_|*z*_*s*_) and *p*_*θ*_(*x*_2_|*z*_*s*_) are calculated with data distribution obtained by the decoders. We estimated the mean joint log-likelihood of the validation set of 100 importance samples (*N*_*s*_ = 100) on all datasets and reported the average values over 5-fold cross-validation.

### 2.5 Evaluation of the cell representations learned by VAEs

We applied the Louvain community detection algorithm [20] to the low-dimensional representations of cells generated by the models on the training and validation sets. Resolution is a parameter that influences the number of identified clusters–a higher value yields more clusters. By running the Louvain clustering over a range of resolution values, we then compare the resulting clusters against the published cell type annotations using the Adjusted Rand Index (ARI) [21] as a measure of how well the learned representations capture the underlying structure of the data.

### 2.6 Data pre-processing

#### Juvenile mouse brain

Bartosovic et al. recently developed nano-CUT&Tag technology, enabling multimodal chromatin profiling at single-cell resolution [1]. The authors succeeded in measuring up to three modalities, ATAC, H3K27ac and H3K27me3, simultaneously within individual cells from the mouse brains (19-day old). Starting with the fragment data of each modality, we used `Signac` to segment the genome into windows, resulting in a count matrix (fragment count in each genomic bin) with the dimension of cell by bin [22]. Fang et al. showed that bin size ranging from 1kb to 10kb performed similarly in their benchmark studies[23]. Therefore, we set the bin width to be 10kb to reduce the input dimension for this analysis. We excluded the bins that overlap with the regions in the ENCODE mouse genome (mm10) blacklist[24]. We retained the cells with the authors’ annotation for the analysis. After filtering, H3K27ac and H3k27me3 were measured in total *N* = 11, 981 cells (4 biological replicates: *N*_1_ = 2, 117, *N*_2_ = 2, 479, *N*_3_ = 2, 392, *N*_4_ = 4, 993), and 4, 434 of them (2 biological replicates: *N*_1_ = 2, 084, *N*_3_ = 2, 350) have additional ATAC measurements. We further selected the 25, 000 bins with the largest counts jointly from all of the modalities (i.e. the union of bins of the highest fragment count of each modality) to reduce sparsity. For bi-modality data, we used 28 cell type labels generated on the H3K27ac (similar to labels generated on H3K27me3) for model performance evaluation. For the dataset encompassing three modalities, we utilized 26 cell classes from WNN analysis conducted by the authors. For convolutional neural networks, we constructed 3-dimensional input arrays (cell *×* modality *×* bin).

#### Human bone marrow mononuclear cells (BMMCs)

Stuart et al. collected bone marrow mononuclear cells from healthy human donors, and jointly profiled H3K27ac and H3K27me3 using single-cell Nanobody-tethered transposition followed by sequencing (scNTT-seq) technology[2]. We downloaded the processed R object from Zenodo (https://zenodo.org/record/7102159), which contains *N* = 5, 236 cells with top 71, 253 bins from H3K27ac modality and top 43, 170 bins from H3K27me3 (bin size = 1 kb) used for the original analysis, where 15 different cell types were identified through WNN workflow on aggregated bin data by the authors. For our analysis, we obtained the genome bin features from fragment files and selected the top 25, 000 bins (bin size = 10 kb) using the same strategy described above.

#### Human peripheral blood mononuclear cells (PBMCs)

The PBMC sample was obtained from a healthy female donor (*N* = 11, 909 before quality control). The dataset was generated by 10x Genomics using single Cell Multiome ATAC + Gene Expression (publicly available on 10x Genomics website). For each cell, 36, 601 genes and 106, 056 peaks were profiled in parallel. We followed the Weighted-Nearest Neighbor (WNN) workflow (Seurat V4) to generate the cell type labels through joint analysis of the transcriptomics (RNA-seq) and chromatin accessibility (ATAC-seq) profiles. We kept the cells (*N* = 11, 402) that meet the specified criteria for quality control (number of ATAC-seq counts *∈* [5, 000, 70, 000]; the number of RNA-seq counts *∈* [1, 000, 25, 000], > 20% mitochondrial counts). Top 5, 000 genes and top 25, 460 peaks were selected for WNN analysis. As a result, the Louvain algorithm (resolution = 0.25) led to 15 clusters, which were further used for method benchmark after cell type annotation. For ATAC peak data, the read (fragment end) count are converted to the fragment count, as suggested by Martens et al. (estimated fragment count = (odd read count + 1)*/*2) [25].

#### Mouse cortex and hippocampus

Yao et al. sequenced approximately 1.3 million cells in the adult mouse cortex and hippocampus regions and obtained their transcriptomic profiles, leading to a thorough assortment of glutamatergic and GABAergic neuron types [26]. For this study, we used the single-cell transcriptomic data generated by 10x Genomics Chromium platform (version 2 chemistry). We downloaded the processed data (*N* = 1, 169, 213) from the Neuroscience Multi-omic (NeMO) Data Archive as part of the BRAIN Initiative Cell Census Network. Out of genes measured in total, we selected 5, 000 highly variable genes from the normalized dataset using LIGER pipeline [27–29]. For evaluation on the investigated methods, we used 42 cell classes and subclasses annotated by the authors following Tasic et al.’s work [30].

#### Mouse organogenesis

Argelaguet et al. investigated the mouse early organogenesis by simultaneously profiling gene expression and chromatin accessibility in the same nuclei (10x Multiome) from mouse embryos between 7.5 to 8.75 days (E7.5-8.75) of gastrulation[31]. Specifically, we selected the E7.5, E8, E8.5 and E8.75 embryos ATAC-seq datasets (*N* = 68, 804), and preprocessed the fragment files following the ArchR pipeline provided by the authors[32]. After excluding the cells identified as low-quality or doublets, we obtained the peak count matrix comprising 191, 407 peaks from *N* = 41, 705 cells. We further selected the 25, 000 peak features with the highest total number of counts across all cells for analysis. Fragment count was estimated using read count following the same approach described above.

### 2.7 Experiments

We benchmarked the selected models through 5-fold cross-validation over a variety of datasets. For model training, we used a mini-batch size of 128, Adam optimizer (learning rate = 0.001). Each fully connected layer has 128 hidden units. The architecture incorporated Batch Normalization and ReLU activation functions in the majority of layers, alongside a dropout rate of 0.2 to prevent overfitting. We performed the Louvain algorithm (*k*-nearest neighbors: *k* = 20) on the latent cell embeddings for clustering. The algorithm was run with 5 random starts unless stated otherwise. The cluster assignment with the best quality was recorded. For the juvenile mouse brain data, the dimension of the latent space *D* was set to 30 and all models underwent 300 epochs of training. For the BMMCs data, we used *D* = 30 and 200 training epochs. For single-cell unimodal datasets: *D* = 20 and 200 training epochs were employed for PBMCs data; *D* = 30 and 15 training epochs for mouse cortex and hippocampus data; *D* = 30 and 50 training epochs for mouse organogenesis data.

### 2.8 Model implementation

All reported VAE models were implemented in Pytorch 1.10.1 and Python 3.8, trained with 2.9 GHz Intel Xeon Gold 6226R and NVIDIA A40 GPU.

## 3 Results

### 3.1 1D-convolutional neural networks for single-cell multimodal epigenomics integration

We introduce `ConvNet-VAE`, a novel approach designed to efficiently learn biologically meaningful low-dimensional cell representations from high-throughput single-cell multimodal epigenomic data. This framework capitalizes on recent advancements in chromatin profiling technologies which permit parallel measurements of histone modifications (e.g., H3K27ac, H3K27me3) and chromatin states at single-cell resolution [1, 2]. The sequenced fragments over the genome are obtained from each individual cell (Figure 1a).

Because single-cell multimodal epigenomic experiments measure different features over the same sequential domain (i.e., the genome), we reasoned that the data is most naturally represented as a multi-channel 1D sequential signal. This is a quite different approach than previous single-cell multimodal neural networks, which treat each modality as if it measured completely unrelated features (e.g., distinct genes or peak locations for each modality). Additionally, previous approaches often use a separate encoder and decoder network for each modality, while ours uses a single encoder and a single decoder that operate on multi-channel signals. By operating on this multi-channel representation of the data, we introduce an appropriate inductive bias that significantly reduces the number of parameters and enforces statistical dependence among neighboring genomic locations within a modality and across modalities at a given genomic locus.

`ConvNet-VAE` is a convolutional variational autoencoder based upon a Bayesian generative model (Figure 1b). To apply 1D-convolutional filters (Conv1D), the input multimodal data are transformed into 3-dimensional arrays (cell *×* modality *×* bin), following window-based genome binning at 10 kilobase resolution [33] (Figure 1a). The encoder efficiently extracts latent factors, which are then mapped back to the input feature space by the decoder network. We use a discrete data likelihood (Poisson distribution) to directly model the observed raw counts.

We also extended `ConvNet-VAE`s to incorporate conditional information such as experimental batches, allowing batch correction using conditional VAEs, which has proven an effective strategy for scRNA-seq data [6]. In our model, the categorical variables (e.g., batch information) are one-hot encoded and then concatenated with the flattened convolutional layer outputs, instead of being combined directly with the multimodal fragment count data over the sorted genomic bins. We incorporated the conditional information in this way because, unlike fully-connected layers, convolutional layers most naturally accommodate sequential data rather than one-hot encodings. Thus, we found it more natural to inject the batch information after the convolutional layers.

In the following sections, we showcase the effectiveness and superiority of `ConvNet-VAE`s by evaluating them on real data and comparing them with `FC-VAE`s.

### 3.2 ConvNet-VAEs learn cell representations using fewer parameters

A key advantage of `ConvNet-VAE`s is the proper inductive bias induced by convolution, which should result in considerable parameter savings. This advantage should increase with the number of modalities per cell: The number of peaks per modality usually exceeds 10^5^, and the peaks often do not overlap across modalities.

To investigate the advantage of `ConvNet-VAE`s on real data, we analyzed a recently published single-cell bi-modal dataset from juvenile mouse brains generated by the nano-CUT&Tag (nano-CT) technology [1]. After preprocessing, the dataset consists of 11, 981 cells from 4 experimental batches with H3K27ac and H3K27me3 modalities. We extracted the top 25, 000 bins identified across both modalities as the input feature set. We then separately examined the effects of (1) kernel size and stride and (2) number of convolutional layers on the number of parameters and performance of `ConvNet-VAE` models. When examining the effects of kernel size and stride, we used architectures with a single convolutional layer and varied kernel size (K) and stride (S) from 11 to 51, with *K* = *S* in each case. Second, we examined the effects of varying the number of convolutional layers from 1 to 3, while keeping a fixed kernel size of 11 and stride of 3. (Note that we used a smaller *S* = 3 with multiple convolutional layers to avoid the output dimensionality being too small.)

We compared all models against `FC-VAE`s. To ensure a fair comparison, we ran all models through 5-fold cross-validation, with 300 training epochs.

Single-Conv1D-layer `ConvNet-VAE`s do indeed require fewer trainable parameters than `FC-VAE`s in this setting. For example, `ConvNet-VAE` (K51, S51) only uses 20% of the parameters that are needed for `FC-VAE`s, while `ConvNet-VAE` (K31, S31) uses 33%. As shown in Figure 2c, as the number of convolutional layers increases, `ConvNet-VAE`s uses fewer parameters. According to the UMAP visualization (colored by the published labels) of the cell embeddings obtained by the selected models, `ConvNet-VAE`s from varying *K, S*, and the number of layers results in qualitatively similar embeddings compared to the `FC-VAE` (Figure 2a).

`ConvNet-VAE`s took slightly longer to complete the training (Figure 2d). The most compact `ConvNet-VAE` (K51, S51) led to a 2.5% decrease in average marginal log-likelihood on the validation sets (Figure 2e), but the (K11, S11) model achieved comparable or better marginal likelihood using 1M fewer parameters than the `FC-VAE`. Increasing the number of convolutional layers or stride resulted in a worse marginal likelihood. However, the models with slightly worse marginal likelihood still excelled in learning low-dimensional cell representations that could reproduce the published cluster assignments (Figure 2f). The Adjusted Rand Index (ARI) first increased as more cell clusters were identified by the Louvain algorithm at a higher clustering resolution, then decreased due to potential over-clustering. `ConvNet-VAE` (K51, S51) achieved the same highest ARI of 0.83 (average over 5 random runs of the Louvain clustering) as single-layer `FC-VAE` did on the training sets, and beat `FC-VAE`s with ARI of 0.82(*±*0.01). Similarly configured `ConvNet-VAE` with smaller kernels and stride displayed a comparable pattern in cluster counts and ARI scores. The two-layer `ConvNet-VAE` performed slightly better in terms of ARI than the one-layer, while one fully-connected layer performed the best, with each additional layer worsening performance. In summary, this first set of tests indicates that `ConvNet-VAE` can achieve similar or better performance compared with `FC-VAE` using fewer parameters.

### 3.3 ConvNet-VAEs show a larger advantage with increasing number of modalities per cell

Because our approach treats each modality as a different channel along a shared sequential domain, we expect the advantage of our approach to increase with the number of modalities profiled per cell. To investigate this, we expanded the analysis by incorporating a 3rd modality, chromatin accessibility, which was measured alongside H3K27ac and H3K27me3 by the developers of nano-CT using assay for transposase-accessible chromatin (ATAC-seq) [1]. A total of 4, 434 cells from 2 experimental batches have ATAC, H3K27ac, and H3K27me3 profiles (three modalities per cell). As in the previous section, we selected the 25, 000 bins with the highest counts across modalities and generated a 4, 434 *×* 3 *×* 25, 000 input for `ConvNet-VAE`s.

Through qualitative evaluation in the UMAP space, the single-Conv1D-layer `ConvNet-VAE` model with a large kernel and stride (K51, S51) results in more compact cell clusters than single-layer `FC-VAE` (Figure 3a), while requiring 87% fewer trainable parameters (Figure 3b). This efficiency remained notable even with a smaller kernel and stride (K11, S11), with a 39% reduction in parameters. The gap in runtime between `ConvNet-VAE`s and `FC-VAE`s also becomes narrower. For instance, single-Conv1D-layer `ConvNet-VAE` (K51, S51) takes 14% more time than the single-layer `FC-VAE`s to finish 300 training epochs, a decrease from the 25% longer runtime seen in the bimodal analysis (Figure 3c). There was no statistical difference in the marginal log-likelihoods across all investigated VAE variants (Figure 3d), implying equivalent capabilities in modeling the data distribution.

**Fig. 3.**
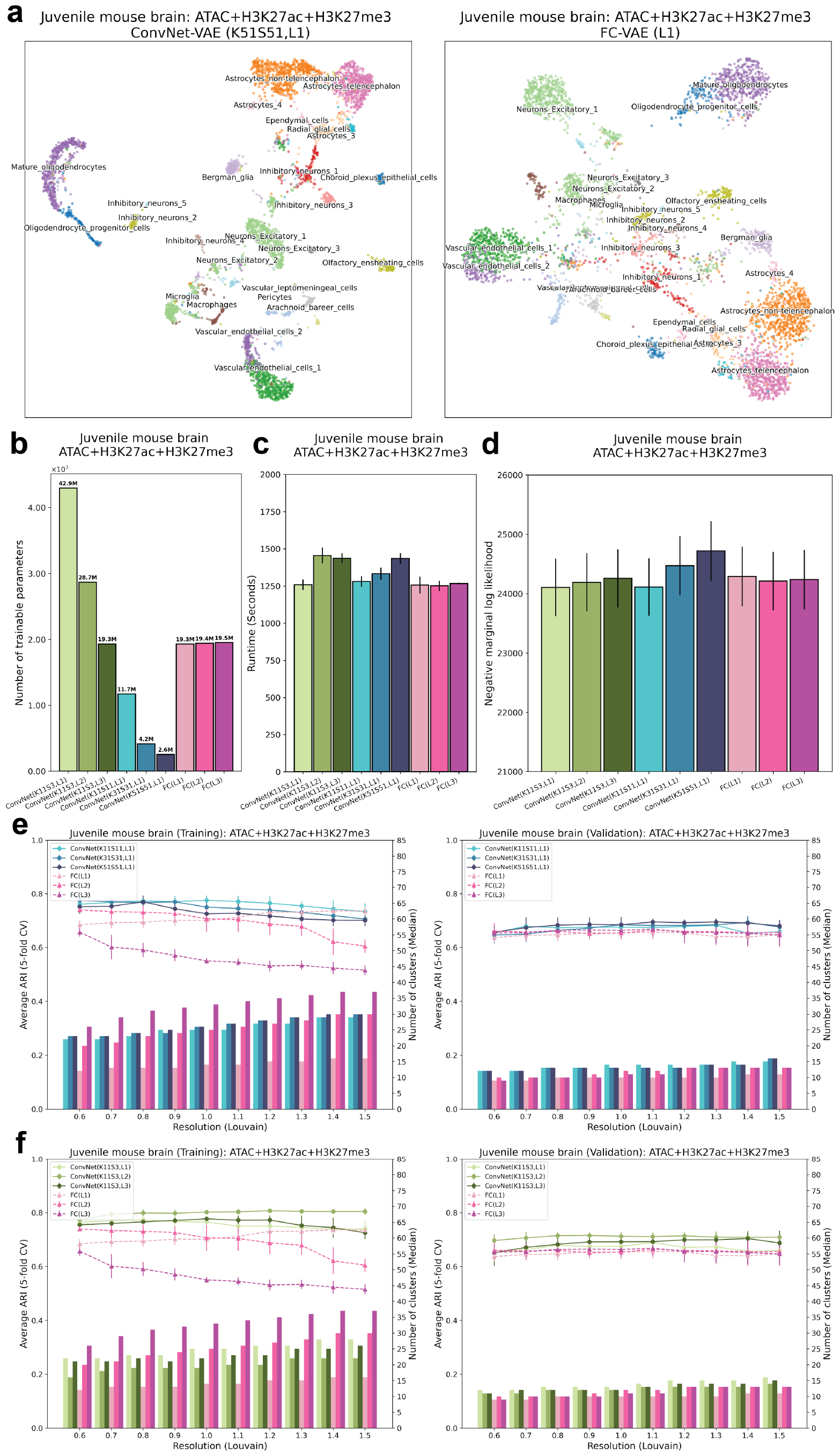
ConvNet-VAEs integrate single-cell tri-modal epigenomic profiling data from mouse brain. (**a**) UMAP visualization of cell embeddings from `ConvNet-VAE`s (Left) and `FC-VAE` (Right). (**b**) The number of trainable parameters of `ConvNet-VAE`s from Group 1 (Blue), Group 2 (Green), and `FC-VAE`s (Pink). (**c**) Average training time is reported for each model. (**d**) Average negative marginal log likelihood of validation set estimated through importance sampling. A lower value implies a larger marginal log likelihood. (**e**,**f**) Comparisons between `ConvNet-VAE`s and `FC-VAE`s in terms of the quality of cell embeddings (training set: left; validation set: right). The bars show the median number of clusters obtained by the Louvain algorithm from 5 splits in cross-validation over a range of resolutions. The corresponding average Adjust Rand Index (ARI) is calculated by comparing the resulting clusters to the published cell type labels, displayed as a line plot. Error bars indicate standard deviation across 5-fold cross-validation.

The advantage of `ConvNet-VAE`s becomes even more apparent when evaluating the quality of the cell embeddings (Figure 3e,f). On both training and validation sets, the single-Conv1D-layer `ConvNet-VAE`s lead in clustering accuracy (highest ARI: 0.78(*±*0.02) at resolution 1.0 for training 0.69(*±*0.01) at resolution 1.1 for validation), as compared to the `FC-VAE`s’ highest ARI of 0.74(*±*0.01) at resolution 0.6 for training and 0.67(*±*0.03) at resolution 1.1 for validation (Figure 3e). This superiority is further supported by the performance of the multi-Conv1D-layer VAEs, which are top performers at almost all clustering resolutions (Figure 3f). For example, 2-Conv1D-layer `ConvNet-VAE` (K11, S3) stands out by producing an ARI of 0.81(*±*0.01) and 0.72(*±*0.01) on the training and validation sets respectively. Interestingly, unlike `FC-VAE`s, where additional layers usually lead to lower quality of the cell latent factors in the training data, `ConvNet-VAE`s can actually benefit from extra convolutional layers (Figure 2g, 3f). Furthermore, the `ConvNet-VAE`s are more effective than `FC-VAE`s as more modalities are added, as evidenced by the collective results from bi-modal and tri-modal integrative analyses.

### 3.4 ConvNet-VAEs allow for improved batch-effect correction

Single-cell data are often generated from different experiments, leading to batch effects that stem from technical rather than biological differences. Therefore, correcting for these effects is essential for clustering and visualization to accurately reflect the underlying biology. A number of methods have been introduced to address this problem in single-cell uni-modal data [17, 27, 28, 34]. The same challenge occurs with these single-cell multimodal epigenomics datasets (Figure 4a, 4c). Without removing batch effects, the cells with bi-modal and tri-modal measurements from different datasets are poorly aligned, resulting in clusters that separate by dataset rather than underlying biological cell type.

**Fig. 4.**
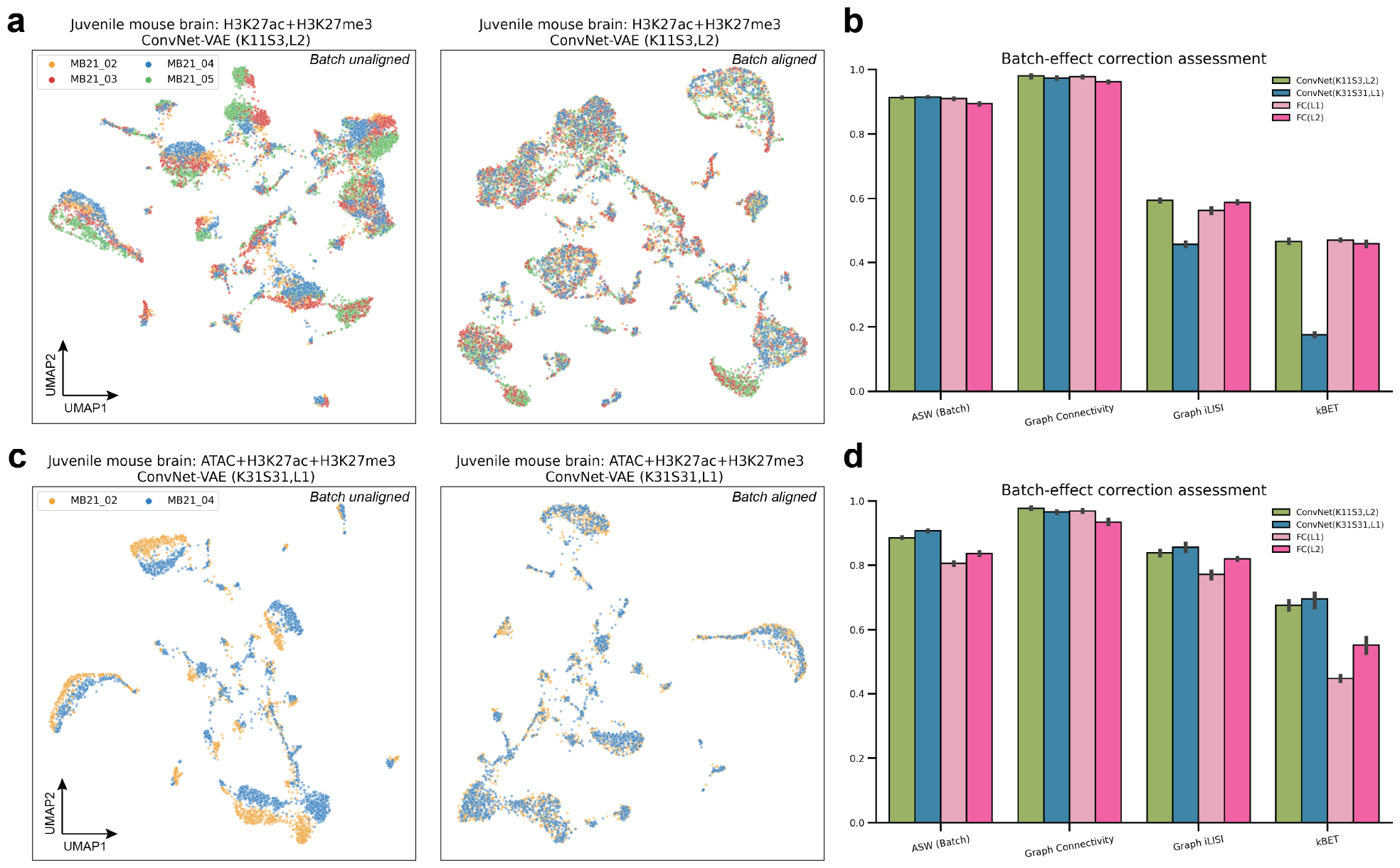
Benchmark of ConvNet-VAEs on batch-effect removal. VAE models were applied on the entire tested datsets without training/validation splitting. (**a, c**) UMAP visualizations of cell embeddings from selected models on the bi- and trimodal data, before and after alignment. (**b, f**) Quantitative comparison between `ConvNet-VAE`s and `FC-VAE`s based on four different metrics. The bars show average scores with standard deviation from 5 random runs.

Here, we selected `ConvNet-VAE`s with single and multiple Conv1D layers to demonstrate their capacity to remove batch-associated technical variation. There are four different batches in the single-cell bi-modal juvenile mouse brain data with measurement of H3K27ac and H3K27me3. When we apply the `ConvNet-VAE` with 2 convolutional layers (K11, S3) with batch information, the cells are well mixed in each cluster (Figure 4a). In the tri-modal setting (simultaneous profiling of chromatin accessibility, H3K27ac, and H3K27me3), the cells from two replicates are clearly separated as shown. Based on the quality of batch mixing, the architecture with a single Conv1D-layer (K31, S31) successfully aligned these cells from both batches (Fig. 4c).

Beyond the qualitative evaluation, we further carried out quantitative assessment of batch-effect removal. To do this, we calculated four metrics: Average silhouette width (ASW, Batch), Graph Connectivity, Graph iLISI, and kBET [16]. In comparison to `FC-VAE`s, both selected `ConvNet-VAE`s showed similar or better performance in terms of ASW (Batch) and Graph connectivity, while the `ConvNet-VAE` with a single convolutional layer (K31, S31) is less favored with respect to Graph iLISI and kBET (Fig. 4b). Encouragingly, the selected `ConvNet-VAE`s excelled across all metrics when more modalities were involved (4d). The improvements in ASW (batch) and k-BET were particularly significant. These results align with the results from the previous section, indicating that `ConvNet-VAE`s show a greater advantage over `FC-VAE`s as the number of epigenomic modalities increases.

### 3.5 ConvNet-VAEs integrate histone modifications from scNTT-seq data

In addition to nano-CT, Stuart et al. [2] developed nanobody-tethered transposition followed by sequencing (scNTT-seq), enabling genome-wide measurement of multiple histone modifications at single-cell resolution.

In this part, we showcase the adaptability and consistent performance of `ConvNet-VAE`s when applied to multimodal data obtained through varied sequencing methods.

Toward this goal, we integrated single-cell bi-modal (H3K27ac and H3K27me3) epigenomic data profiled from bone marrow mononuclear cells (BMMCs) of healthy human donors (*N* = 5, 236). According to the UMAP plots, H3K27ac itself doesn’t carry sufficient information to distinguish different cell types, whereas H3K27me3 provides sufficient information to identify the major cell types. Combining both modalities with the selected single-layer `ConvNet-VAE` (K11, S11), we achieved more compact cell clusters (Figure 5a).

**Fig. 5.**
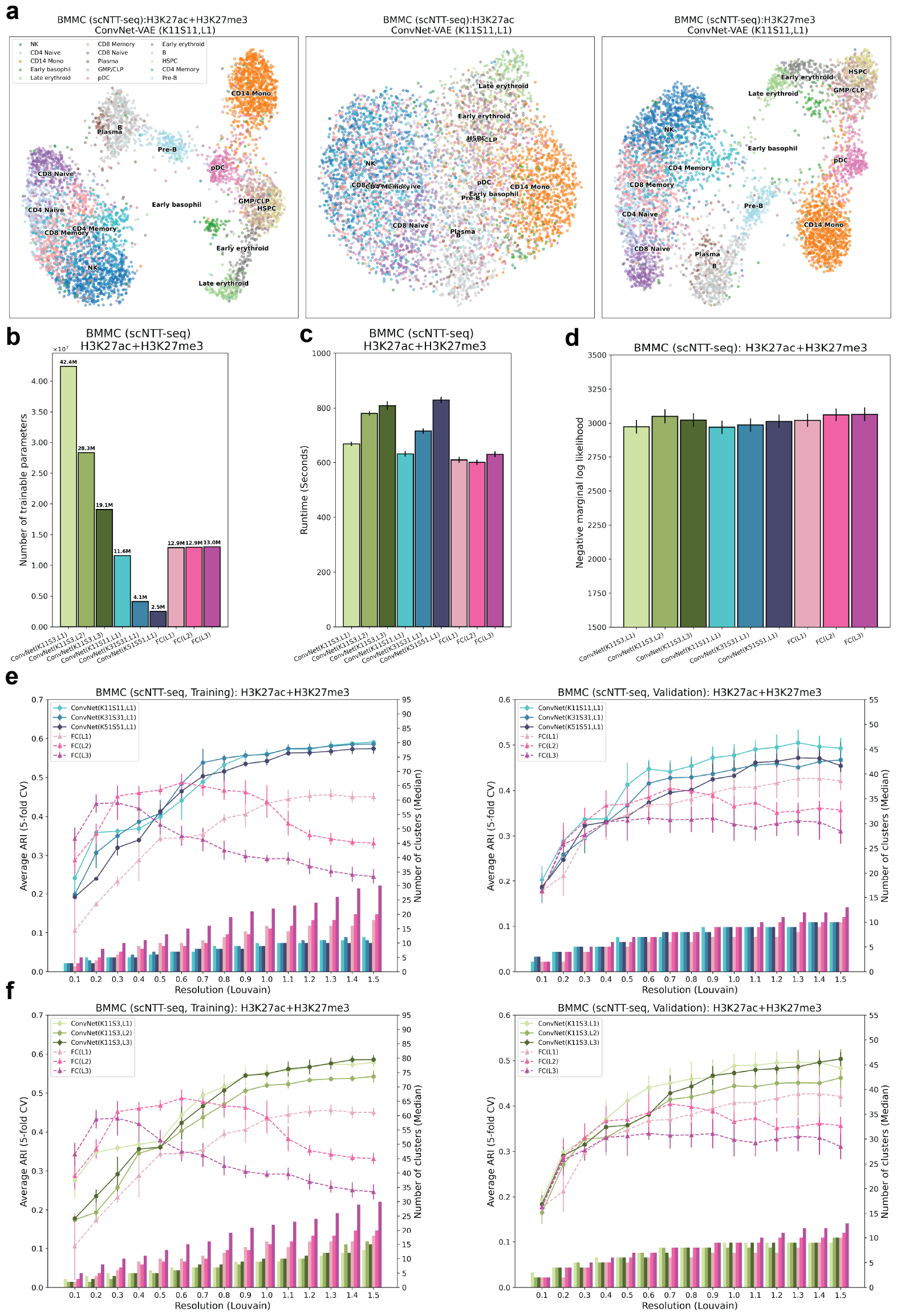
ConvNet-VAEs effectively integrate scNTT-seq data. (**a**) UMAP visualization of cell embeddings from `ConvNet-VAE` (single Conv1D layer, Kernel size 11, stride 11) on BMMCs data containing H3K27ac and H3K27me3 (Left), H3K27ac (Middle), and H3K27me3 (Right). (**b**) The number of trainable parameters of `ConvNet-VAE`s from Group 1 (Blue), Group 2 (Green), and `FC-VAE`s (Pink). (**c**) Average training time is reported for each model. (**d**) Average negative marginal log likelihood of validation set estimated through importance sampling. A lower value implies a larger marginal log likelihood. (**e**,**f**) Comparisons between `ConvNet-VAE`s and `FC-VAE`s in terms of the quality of cell embeddings (training set: left; validation set: right). The bars show the median number of clusters obtained by the Louvain algorithm from 5 splits in cross-validation over a range of resolutions. The corresponding average Adjust Rand Index (ARI) is calculated by comparing the resulting clusters to the published cell type labels, displayed as a line plot. Error bars indicate standard deviation across 5-fold cross-validation.

Although training `ConvNet-VAE`s with multiple convolutional layers, or those with larger kernels and strides, might require additional time compared to `FC-VAE`s (Figure 5c), the `ConvNet-VAE`s display comparable or superior performance in estimating the marginal log-likelihood for the validation data, making them preferable as generative models (Figure 5d). More strikingly, the proposed `ConvNet-VAE`s outperform the `FC-VAE`s on the training and validation sets by a large margin, when comparing ARI as the measure of the effectiveness of dimension reduction (Figure 5e,f).

In order to examine whether the convolutional layers are able to exploit the sequential relationships among genomic locations, we randomly shuffled genomic bins from this BMMC dataset and re-analyzed it with `ConvNet-VAE`s. The decline in ARI upon bin shuffling confirmed that the convolutional layers are indeed sensitive to local epigenomic patterns. In terms of the marginal log-likelihood, the negative effect brought by bin shuffling becomes more apparent when a larger kernel size is used (Supplementary Figure S6). All these observations underscore the ability of 1D-convolutional layers to capture spatial dependencies in the tested single-cell multimodal epigenomic data.

Moreover, we investigated the applicability of `ConvNet-VAE`s on unimodal single-cell data. In analyses of PBMCs gene expression, PBMC ATAC (peaks), as well as the mouse organogenesis ATAC (peaks), single-Conv1D-layer `ConvNet-VAE`s perform on par with `FC-VAE`s. `ConvNet-VAE`s lead the performance in reducing the dimension of the large-scale mouse cortex and hippocampus transcriptomic profile (Supplementary Figures S7,S8,S9,S10).

## 4 Discussion

In this study, we proposed the `ConvNet-VAE` framework, specifically designed to model single-cell multimodal epigenomic data. This model comprises 1D-convolutional layers and hence takes multi-channel binned fragment counts as input. The encoder network within this framework learns low-dimensional representations of cells that facilitate cell type inference following clustering. We validated `ConvNet-VAE`s’ utilities through integrative analyses of bi-modal (H3K27ac + H3K27me3) and tri-modal juvenile mouse brain data (ATAC + H3K27ac + H3K27me3), as well as bi-modal data from human bone marrow mononuclear cells (H3K27ac + H3K27me3).

As demonstrated by the results, `ConvNet-VAE`s are able to extract information about chromatin states and histone modifications, accurately capture the data distribution, and correct for batch effects. The 1D-convolution layers are capable of capturing the spatial relationships among sequentially arranged genomic bins. In qualitative and quantitative benchmarking with `FC-VAE`s, which solely utilize fully connected layers, `ConvNet-VAE`s show effectiveness by achieving on-par or enhanced performance using far fewer parameters. Unlike `FC-VAE`s, `ConvNet-VAE`s can also benefit from including more layers (Conv1D) in the model architecture. Notably, the advantage of `ConvNet-VAE`s over `FC-VAE`s becomes more evident when jointly analyzing three modalities instead of two.

Nevertheless, the `ConvNet-VAE`s presented in this report are not without limitations. Due to the use of convolutional filters, they require that all modalities share the same feature space (i.e. an identical set of bins). Moreover, there is potential to further refine model performance by optimizing parameters like kernel size and stride length.

To summarize, the `ConvNet-VAE` framework stands out for its performance in integrating single-cell multimodal epignomic data. We anticipate that the utility of our approach will become more promising as the number of modalities and cells in single-cell multimodal epigenomic datasets increases in the future.

## 5 Data Availability

The published raw datasets are listed below. The processed count data used for analyses can be accessed on figshare (https://figshare.com/s/e5696dd20f7b0603f122).

- Juvenile mouse brain from Bartosovic et al. (2023) (GSE198467)
- Human bone marrow mononuclear cells (BMMCs) from Stuart et al. (2022) (GSE212588)
- Human peripheral blood mononuclear cells from 10x Genomics (2021) (https://www.10xgenomics.com/resources/datasets)
- Mouse cortex and hippocampus from Yao et al. (2021) (https://data.nemoarchive.org/biccn/grant/u19_zeng/zeng/transcriptome/scell/10x_v2/mouse/processed/YaoHippo2020/)
- Mouse organogenesis from Argelaguet et al. (2022) (GSE205117)

The source code of VAE models and a demo in Jupyter notebook are provided in GitHub repository (https://github.com/welch-lab/ConvNetVAE).

## 6 Acknowledgements

The authors thank Matthew Karikomi and Jie Liu for helpful discussions. This work was supported by NIH grant R01 HG010883 to J.D.W.

## 8 Author Contributions

JDW and CG conceived the idea of `ConvNet-VAE`. CG developed and implemented the model. CG conducted the data analyses. CG and JDW wrote the paper. All authors read and approved the final manuscript.

## Supplementary Figures

**Fig. S1.**
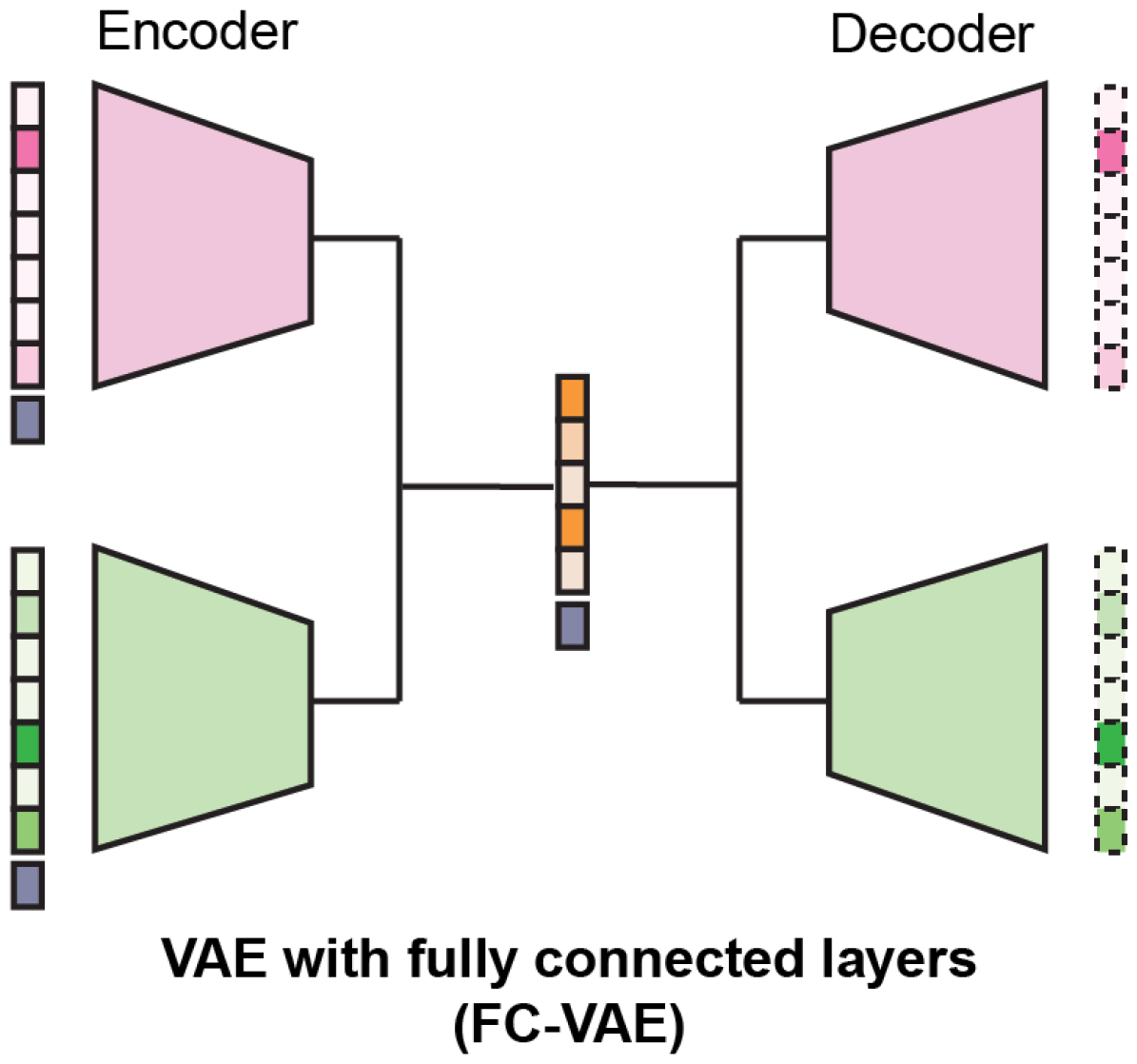
Architecture of FC-VAE (bi-modal). A brief illustration of the architecture of a bi-modal `FC-VAE` based on Product of Experts (PoE). Each expert corresponds to a modality. It easily extends to additional modalities by adding encoder-decoder pairs.

**Fig. S2.**
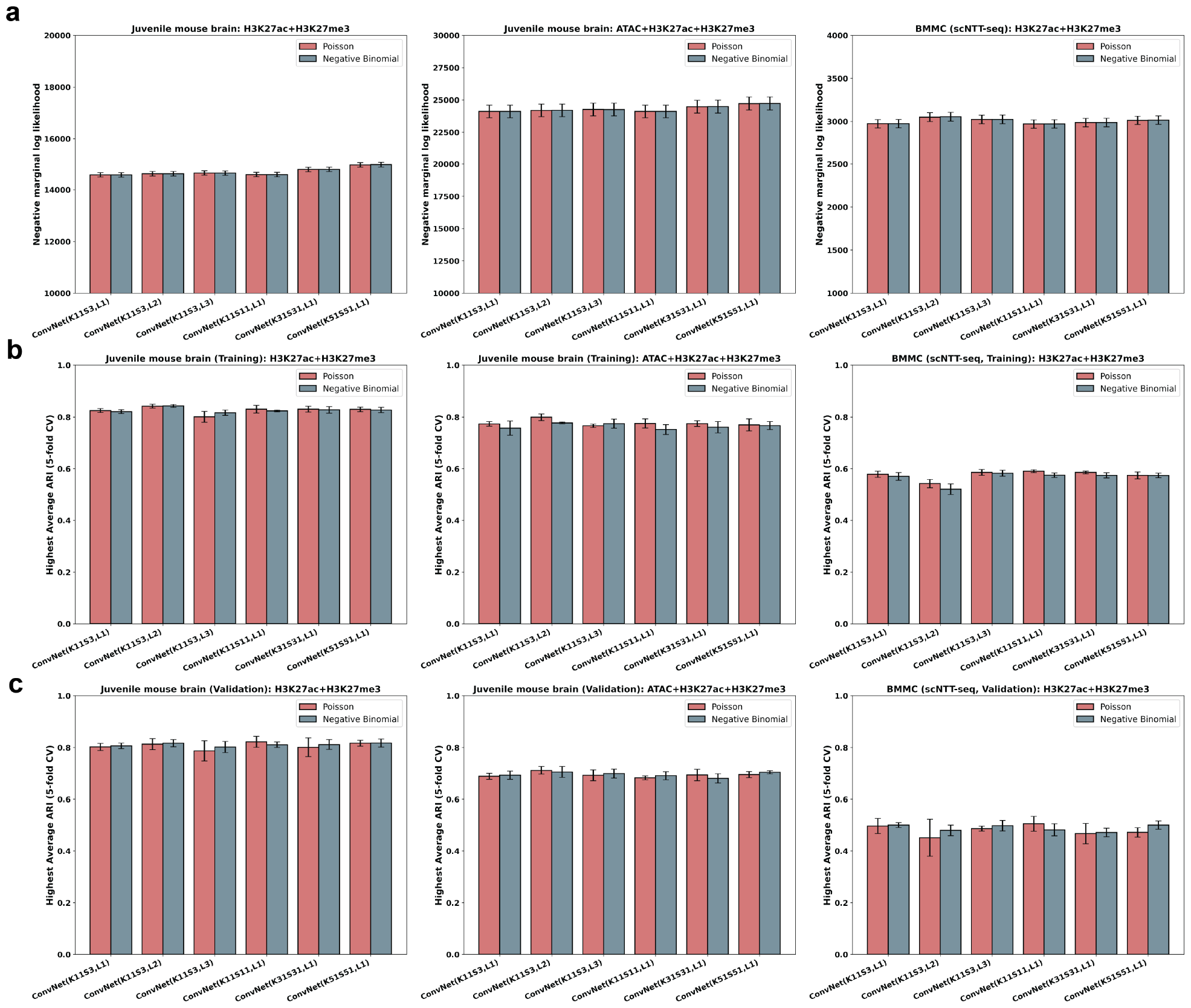
Models with Poisson and negative binomial distributions lead to comparable performance on studied datasets. Bi-modal juvenile mouse brain (Left column), tri-modal juvenile mouse brain (Middle column), BMMCs (Right column). (**a**) Comparison of the marginal log likelihood (validation set) from `ConvNet-VAE`s under Poisson and negative binomial distributional assumption. (**b**) The highest average ARI that each model can achieve on the training sets over a range of clustering resolutions. (**c**) The highest average ARI that each model can achieve on the validation sets over a range of clustering resolutions. All error bars indicate the standard deviation from 5-fold cross-validation.

**Fig. S3.**
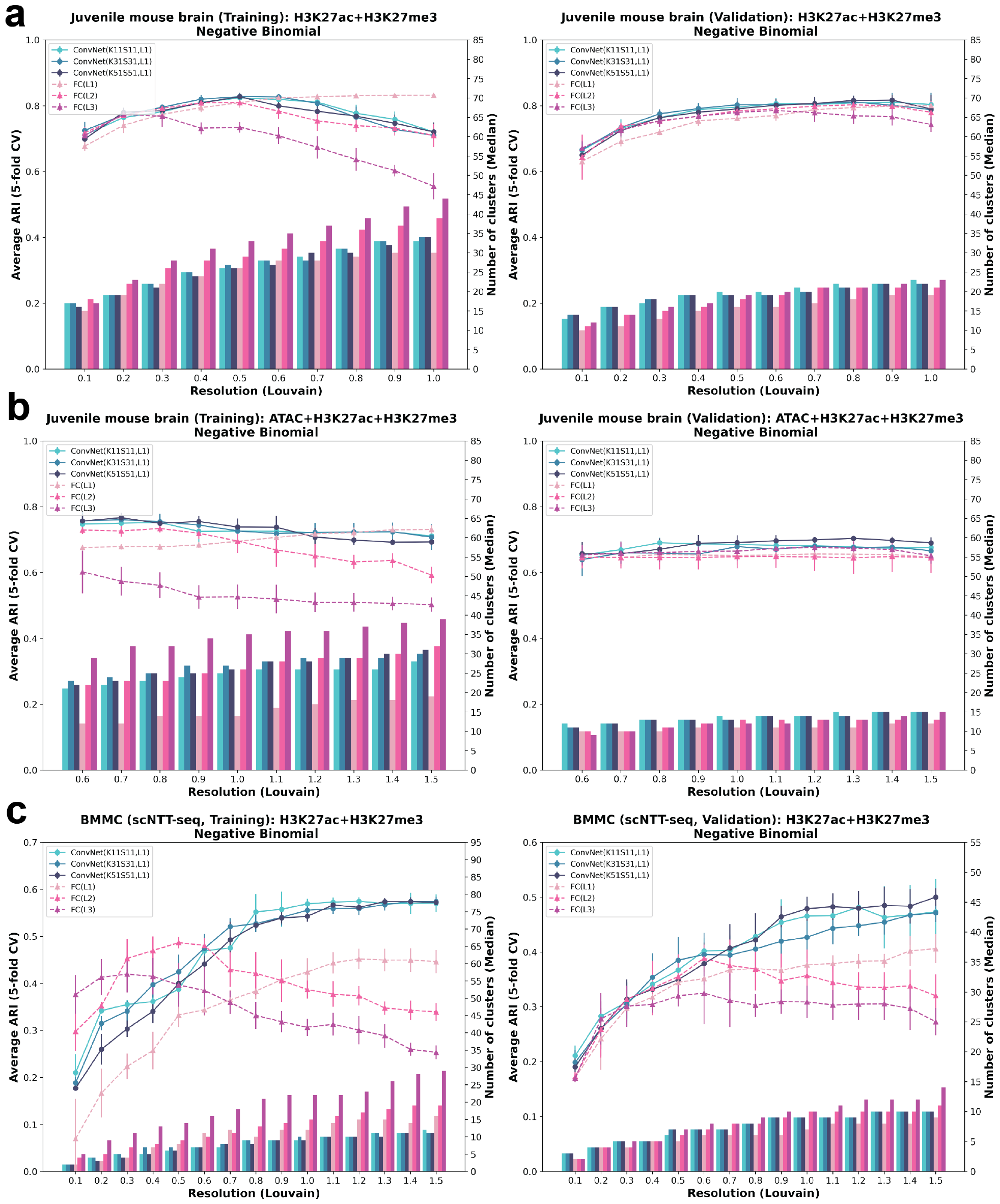
Evaluation of 1-Conv1D-layer ConvNet-VAEs with negative binomial distribution: ARI. Comparison between `ConvNet-VAE`s (Group 1) using negative binomial modeling and `FC-VAE`s on the quality of cell embeddings, evaluated by ARI. (**a**) Bi-modal juvenile mouse brain. (**b**) Tri-modal juvenile mouse brain. (**c**) BMMCs. Error bars indicate the standard deviation from 5-fold cross-validation.

**Fig. S4.**
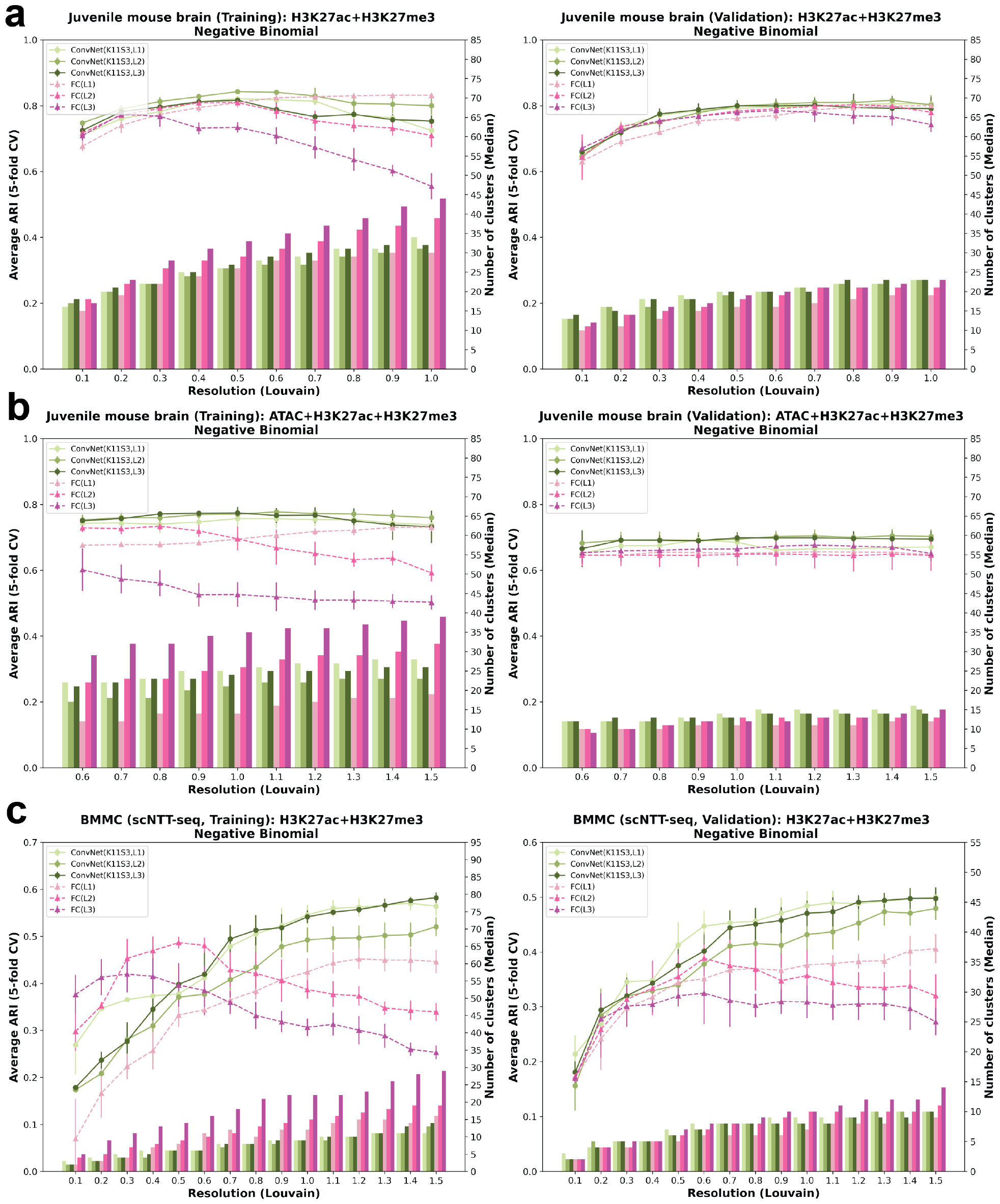
Evaluation of multi-Conv1D-layer ConvNet-VAEs with negative binomial distribution: ARI. Comparison between `ConvNet-VAE`s (Group 2) using negative binomial modeling and `FC-VAE`s on the quality of cell embeddings, evaluated by ARI. (**a**) Bi-modal juvenile mouse brain. (**b**) Tri-modal juvenile mouse brain. (**c**) BMMCs. Error bars indicate the standard deviation from 5-fold cross-validation.

**Fig. S5.**
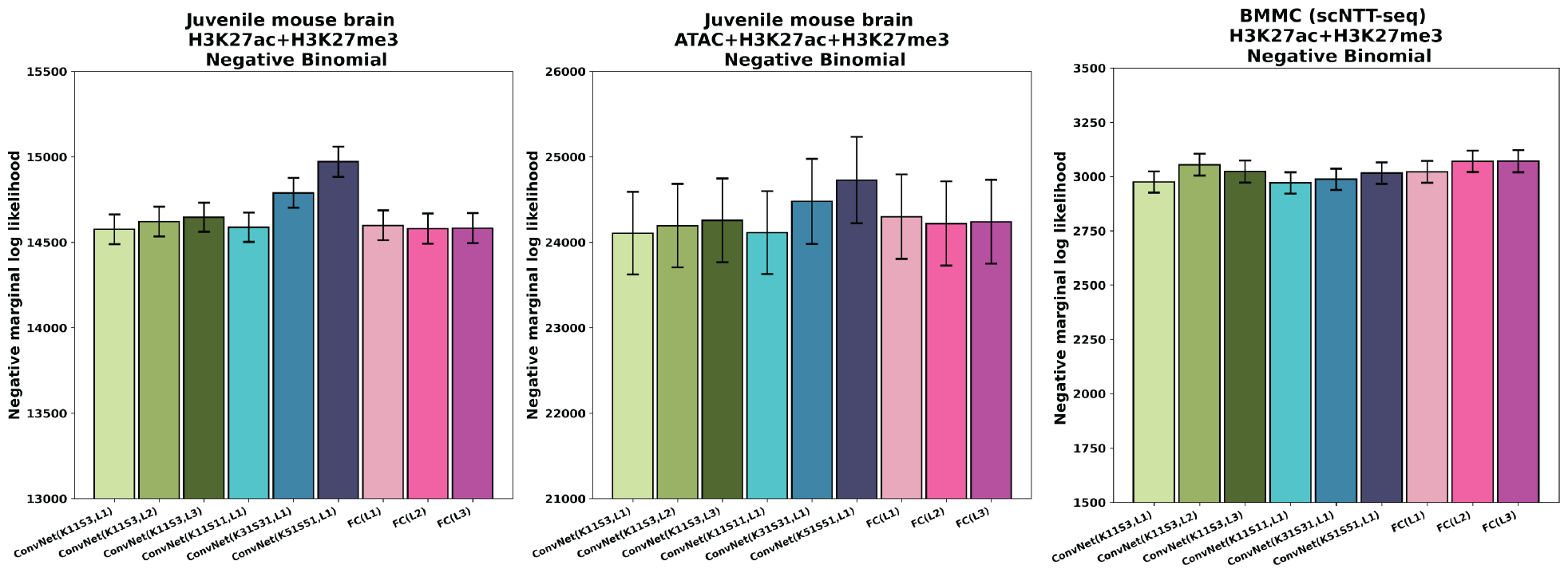
Evaluation of ConvNet-VAEs with negative binomial: Marginal log likelihood (Validation). Comparison of the marginal log likelihood of the validation set between `ConvNet-VAE`s (negative binomial modeling) and `FC-VAE`s.

**Fig. S6.**
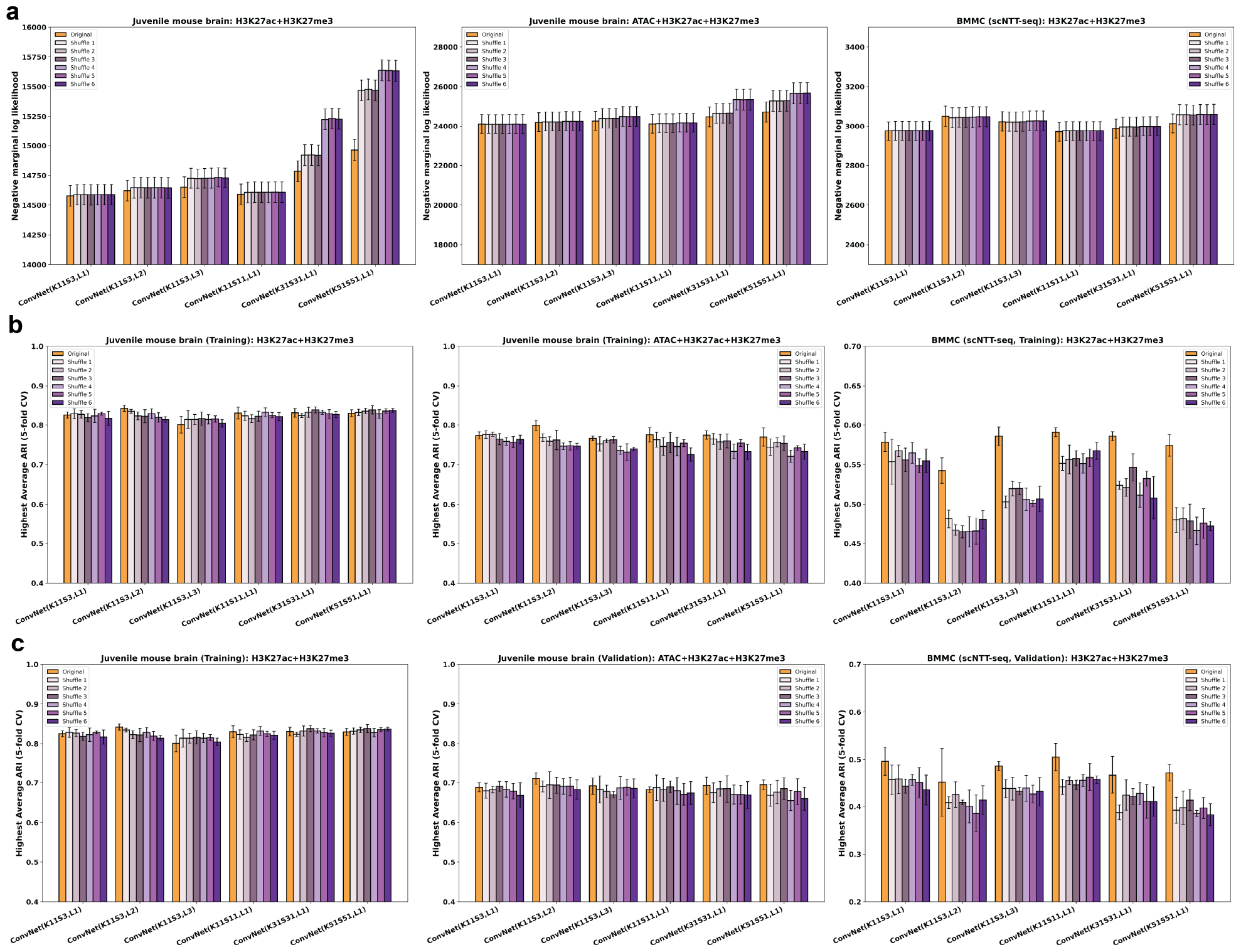
Performance of ConvNet-VAEs after shuffling genomic bins. Side-by-side comparison between the results from the ordered bins and shuffled bins. Bi-modal juvenile mouse brain (Left column), tri-modal juvenile mouse brain (Middle column), BMMCs (Right column). (**a**) Comparison of the marginal log likelihood (validation set). (**b**) The highest average ARI that each model can achieve on the training sets over a range of clustering resolutions. (**c**) The highest average ARI that each model can achieve on the validation sets over a range of clustering resolutions. Error bars indicate the standard deviation from 5-fold cross-validation.

**Fig. S7.**
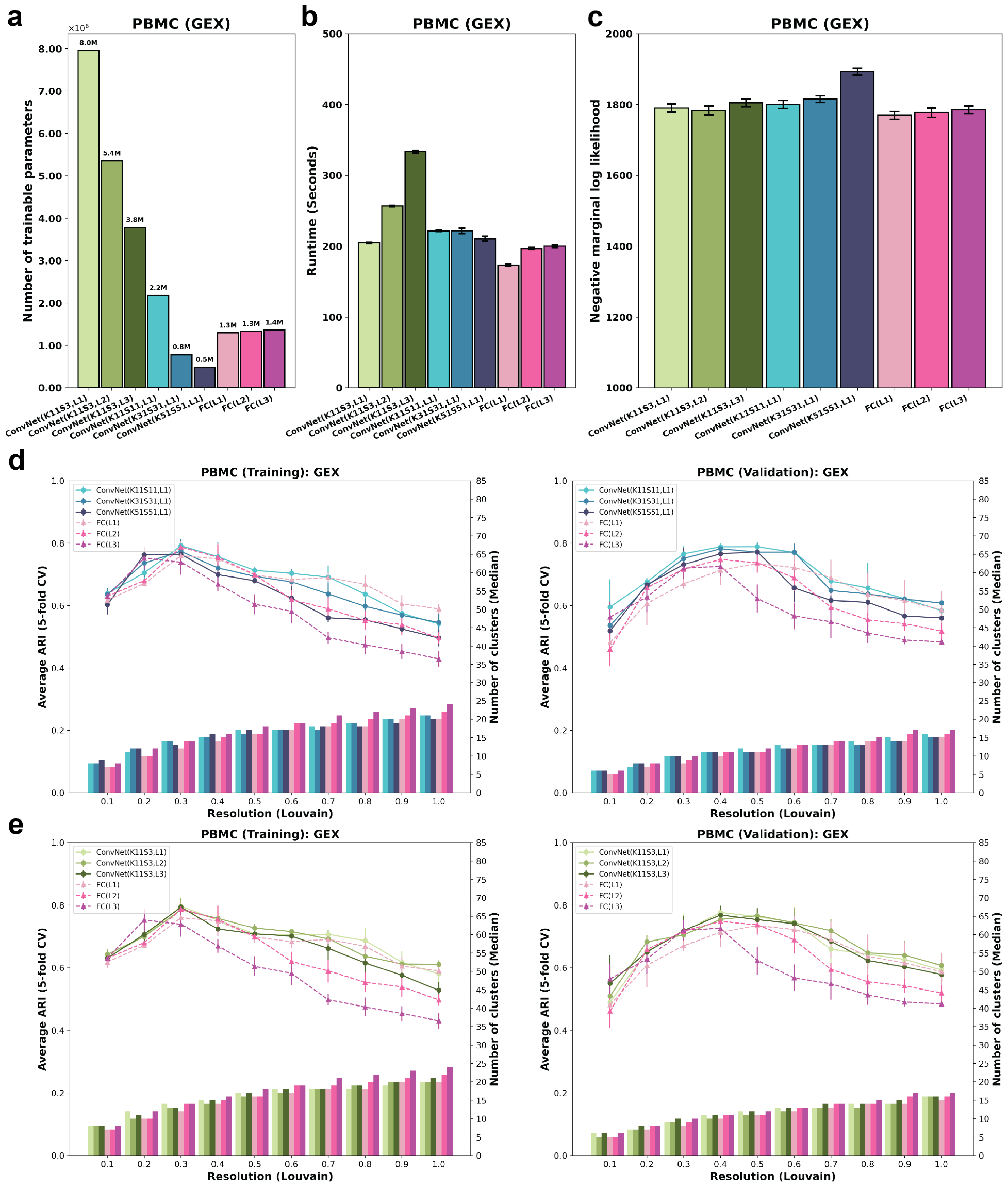
Evaluation of ConvNet-VAEs on PBMCs (gene expression). (**a**) The number of trainable parameters of `ConvNet-VAE`s from Group 1 (Blue), Group 2 (Green), and `FC-VAE`s (Pink). (**b**) Average training time is reported for each model. Error bars indicate standard deviation across 5-fold cross-validation. (**c**) Average negative marginal log likelihood of validation set estimated through importance sampling. (**d**,**e**) Comparisons between `ConvNet-VAE`s and `FC-VAE`s on cell embeddings’ quality. The bars show the median number of clusters obtained by the Louvain algorithm from 5 splits in cross-validation over a range of resolutions. The corresponding average Adjust Rand Index (ARI) is calculated by comparing the resulting clusters to the published cell type labels (line plot). Error bars indicate standard deviation across 5-fold cross-validation.

**Fig. S8.**
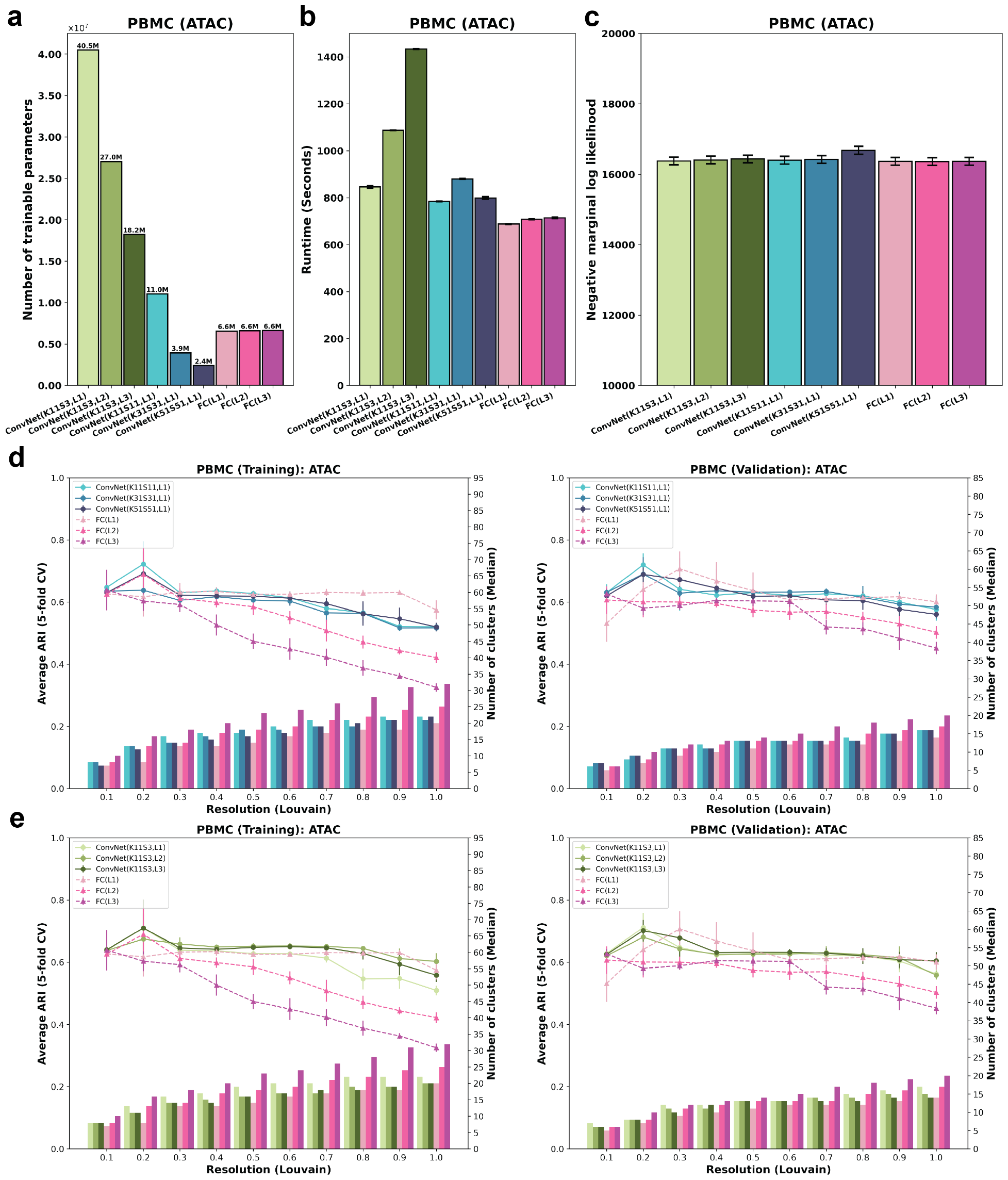
Evaluation of ConvNet-VAEs on PBMCs (ATAC peaks). (**a**) The number of trainable parameters of `ConvNet-VAE`s from Group 1 (Blue), Group 2 (Green), and `FC-VAE`s (Pink). (**b**) Average training time is reported for each model. Error bars indicate standard deviation across 5-fold cross-validation. (**c**) Average negative marginal log likelihood of validation set estimated through importance sampling. (**d**,**e**) Comparisons between `ConvNet-VAE`s and `FC-VAE`s on cell embeddings’ quality. The bars show the median number of clusters obtained by the Louvain algorithm from 5 splits in cross-validation over a range of resolutions. The corresponding average Adjust Rand Index (ARI) is calculated by comparing the resulting clusters to the published cell type labels (line plot). Error bars indicate standard deviation across 5-fold cross-validation.

**Fig. S9.**
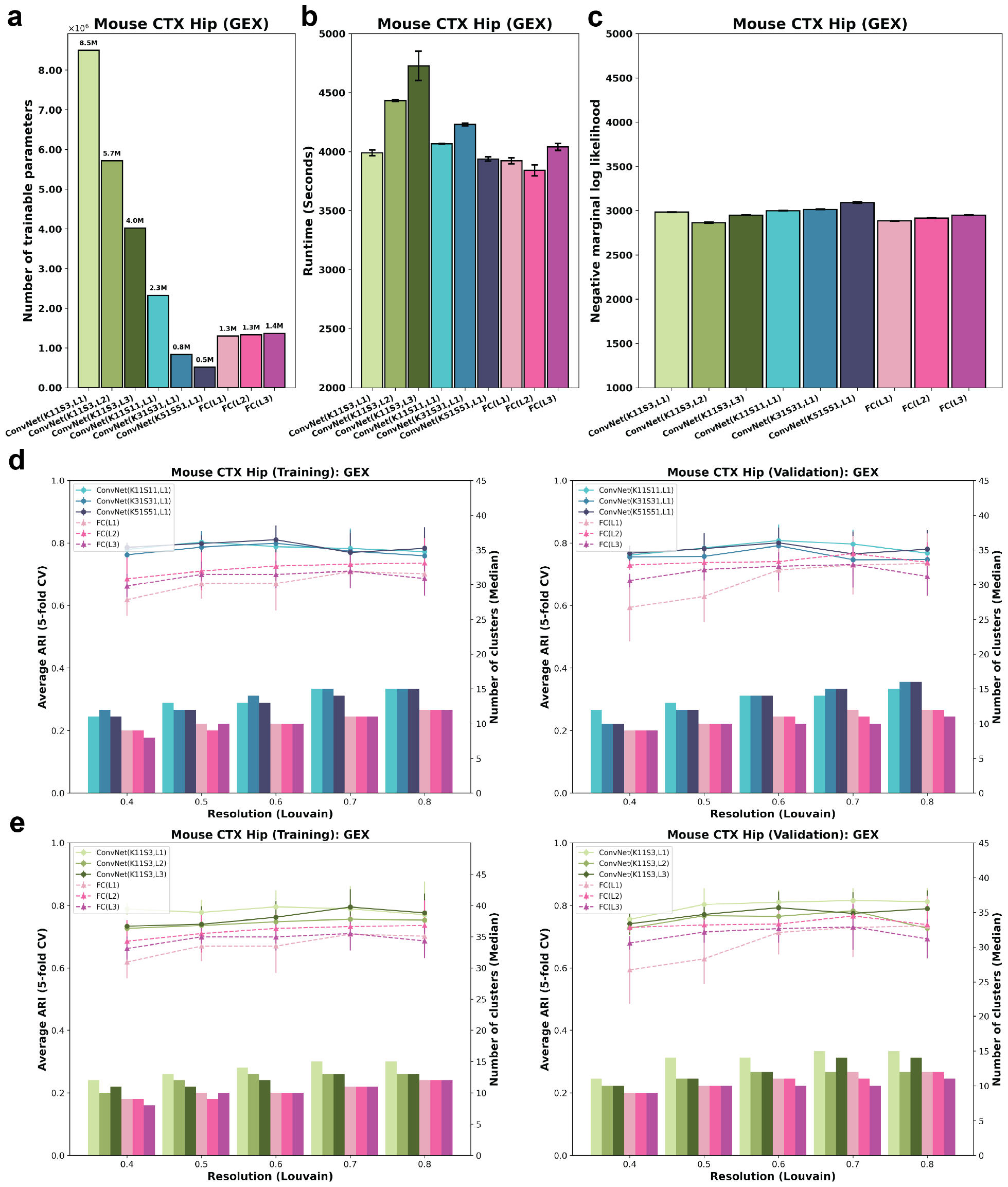
Evaluation of ConvNet-VAEs on mouse cortex and hippocampus (gene expression). (**a**) The number of trainable parameters of `ConvNet-VAE`s from Group 1 (Blue), Group 2 (Green), and `FC-VAE`s (Pink). (**b**) Average training time is reported for each model. Error bars indicate standard deviation across 5-fold cross-validation. (**c**) Average negative marginal log likelihood of validation set estimated through importance sampling. (**d**,**e**) Comparisons between `ConvNet-VAE`s and `FC-VAE`s on cell embeddings’ quality. The bars show the median number of clusters obtained by the Louvain algorithm from 5 splits in cross-validation over a range of resolutions. The corresponding average Adjust Rand Index (ARI) is calculated by comparing the resulting clusters to the published cell type labels (line plot). Error bars indicate standard deviation across 5-fold cross-validation.

**Fig. S10.**
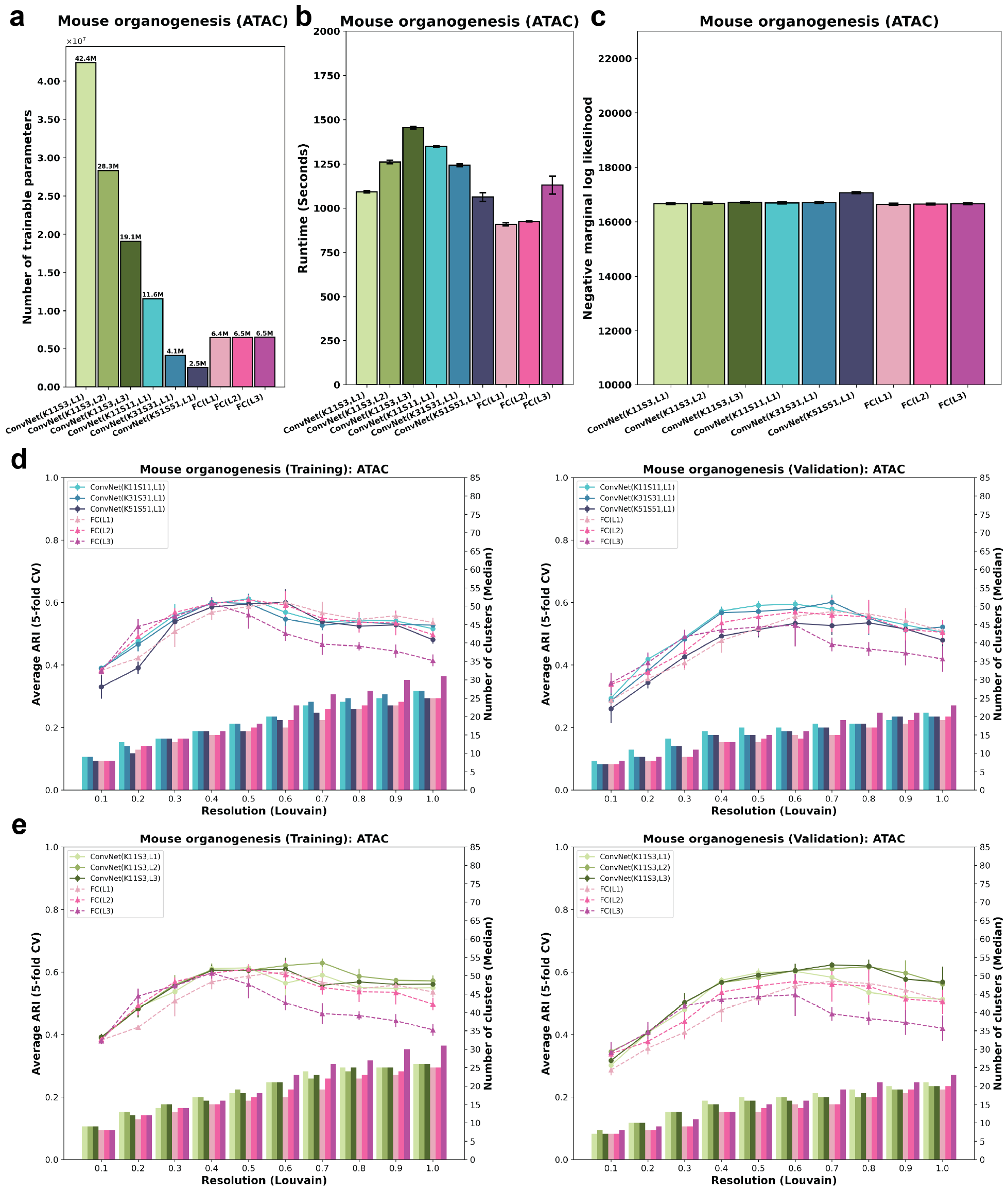
Evaluation of ConvNet-VAEs on mouse organogenesis (ATAC peaks). (**a**) The number of trainable parameters of `ConvNet-VAE`s from Group 1 (Blue), Group 2 (Green), and `FC-VAE`s (Pink). (**b**) Average training time is reported for each model. Error bars indicate standard deviation across 5-fold cross-validation. (**c**) Average negative marginal log likelihood of validation set estimated through importance sampling. (**d**,**e**) Comparisons between `ConvNet-VAE`s and `FC-VAE`s on cell embeddings’ quality. The bars show the median number of clusters obtained by the Louvain algorithm from 5 splits in cross-validation over a range of resolutions. The corresponding average Adjust Rand Index (ARI) is calculated by comparing the resulting clusters to the published cell type labels (line plot). Error bars indicate standard deviation across 5-fold cross-validation.

